# Indirect genetic effects increase the heritable variation available to selection and are largest for behaviours: a meta-analysis

**DOI:** 10.1101/2024.05.17.594196

**Authors:** Francesca Santostefano, Maria Moiron, Alfredo Sánchez-Tójar, David N Fisher

**Affiliations:** University of Exeter, Centre for Ecology and Conservation, University of Exeter, Penryn Campus, Cornwall, U.K.; Université du Québec à Montréal, Département des Sciences Biologiques, Montréal, Canada; Institute of Avian Research, Wilhelmshaven, 26386, Germany; Department of Evolutionary Biology, Bielefeld University, 33615, Bielefeld, Germanyc; School of Biological Sciences, University of Aberdeen, King’s College, Aberdeen, United Kingdom

## Abstract

The evolutionary potential of traits is governed by the amount of heritable variation available to selection. While this is typically quantified based on genetic variation in a focal individual for its own traits (direct genetic effects, DGEs), when social interactions occur, genetic variation in interacting partners can influence a focal individual’s traits (indirect genetic effects, IGEs). Theory and studies on domesticated species have suggested IGEs can greatly impact evolutionary trajectories, but whether this is true more broadly remains unclear. Here we perform a systematic review and meta-analysis to quantify the amount of trait variance explained by IGEs and the contribution of IGEs to predictions of adaptive potential. We identified 180 effect sizes from 47 studies across 21 species and found that, on average, IGEs of a single social partner account for a small but statistically significant amount of phenotypic variation (0.03). As IGEs affect the trait values of each interacting group member and due to a typically positive – although statistically nonsignificant – correlation with DGEs (*r*_DGE-IGE_ = 0.26), IGEs ultimately increase trait heritability substantially from 0.27 (narrow-sense heritability) to 0.45 (total heritable variance). This 66% average increase in heritability suggests IGEs can increase the amount of genetic variation available to selection. Furthermore, whilst showing considerable variation across studies, IGEs were most prominent for behaviours, and to a lesser extent for reproduction and survival, in contrast to morphological, metabolic, physiological, and development traits. Our meta-analysis therefore shows that IGEs tend to enhance the evolutionary potential of traits, especially for those tightly related to interactions with other individuals such as behaviour and reproduction.

**Lay Summary:** Predicting evolutionary change is important for breeding better livestock and crops, for understanding how biodiversity arises and how populations respond to environmental change. Normally, these predictions are based on how the genetic variants in an organism influence its own traits (characteristics). However, when organisms socially interact, for instance by fighting or cooperating, then the genes in one individual can influence the traits of others, therefore affecting the potential for evolutionary change. We compared 47 studies across 21 animal species and found that the effect of the genes of a single social partner is small but statistically significant, while the total contribution of social genetic effects to evolutionary potential is large. These effects are particularly important for the evolution of animals’ behaviours and reproductive traits, but less so for other traits such as body size and physiology. We also found that, because an individual can interact with many others and influence them all, social interactions can substantially increase the potential for a population to evolve from generation to generation. Our results show how social interactions can potentially alter the evolution of those traits known to respond to social interactions in comparison to standard expectations.

## Introduction

Traits can evolve when they are underpinned by heritable genetic variation and subject to selection. The (narrow-sense) heritability of a trait is defined as the ratio of the additive genetic variance to the total phenotypic variance within a population, with a higher ratio indicating greater potential for evolutionary change in response to selection (Falconer & Mackay, 1996; Fisher, 1930; Lynch & Walsch, 1998). Environmental variation generally makes up the majority of the remainder of the total phenotypic variation and is thought not to contribute to evolved changes. However, the environment of an individual can also include its conspecifics and their traits, such as the phenotype of the breeding partner, a neighbouring individual, or a competitor. Most organisms live at least part of their lives socially, interacting when they mate, fight, cooperate, and compete for access to resources (Frank, 2007; Székely et al., 2010; Westneat & Fox, 2010). Through these social interactions, individuals can influence the phenotypic expression of traits in conspecifics. If an individual’s effect on others has a heritable component, then genes in one individual influence the phenotype of others, known as indirect genetic effects (hereafter IGEs; Fig. 1; Dickerson, 1947; Griffing, 1967; Moore et al., 1997; Scott & Fuller, 1965; Wolf et al., 1998). These IGEs can contribute to phenotypic evolution alongside the additive genetic variance stemming from an organism’s own genes (direct genetic effects; hereafter DGEs; Fig. 1). However, when measuring the evolutionary potential of traits, researchers typically only quantify DGEs. Only considering DGEs neglects the total genetic variance in trait values stemming from other sources, such as IGEs, and can lead to inaccurate evolutionary predictions (Wolf et al., 1998).

**Figure 1.**
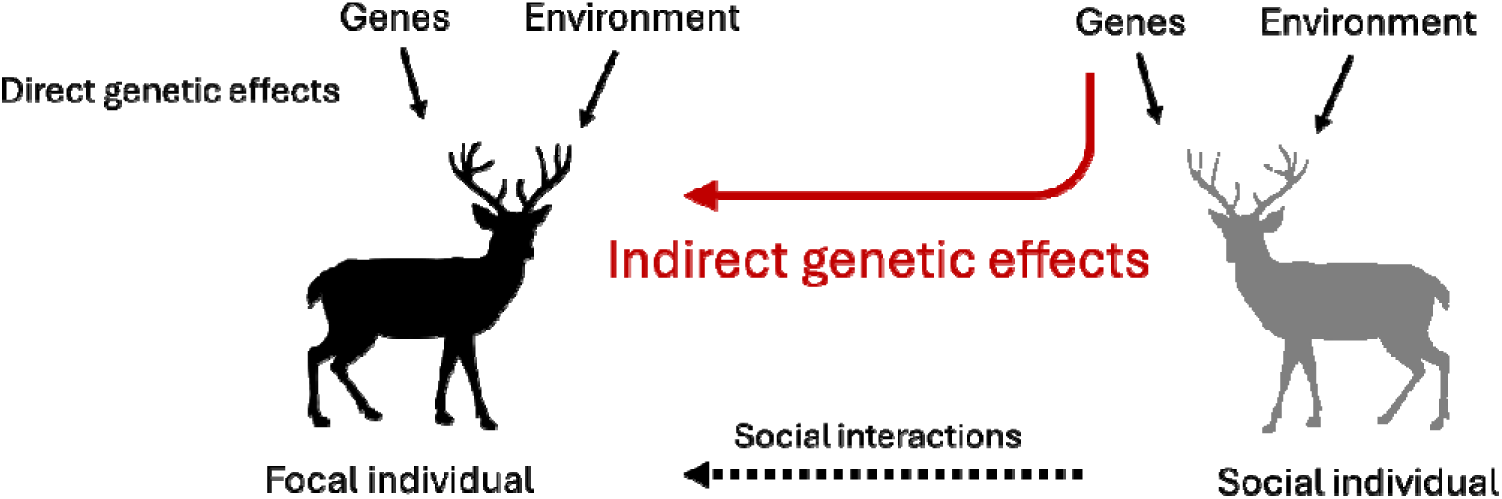
Indirect genetic effects occur when the genotype of one individual influences the expression of a phenotype in a conspecific, mediated by social interactions.

Although the extra genetic variance generated by IGEs can speed up evolutionary change, IGEs can also reduce, remove, or even reverse the response to selection if DGEs and IGEs are negatively correlated (Griffing, 1967; Moore et al., 1997). A positive DGE-IGE correlation indicates that individuals with genes for high trait values elicit higher trait values in their partners (e.g. aggressive individuals instigating higher aggression in others, Alemu et al., 2014; Wilson et al., 2009). In contrast, a negative correlation indicates individuals with high trait values are associated with low trait values in partners (e.g. individuals good at acquiring limited resources cause others to acquire fewer resources, Wilson, 2014). IGEs have long been appreciated in livestock breeding, where they can be exploited to increase productivity and improve welfare (see Ellen et al., 2014 for a qualitative review), as well as in arable farming to increase yield through intercropping (Wright 1985, Bourke et al. 2021), and recently in human medicine (Baud et al. 2017). IGEs have also been studied in the context of maternal genetic effects (the heritable effect a mother has on the traits of her offspring beyond the direct effect of the genes they inherit from her; Dickerson, 1947; McAdam et al., 2014; Mousseau & Fox, 1998), as well as implicitly within social and sexual runaway dynamics (Bailey & Kölliker, 2019). Recently, however, we have begun to appreciate that IGEs are applicable to a much broader array of traits and contexts (McAdam et al., 2014). Examples include the influence of a male bird on the lay date of his mate (Brommer & Rattiste, 2008; Evans et al., 2020), the elicitation of aggression from an opponent during a contest (Santostefano et al., 2017; Wilson et al., 2009), and the growth rate and infection status of neighbours (Costa E Silva et al., 2013). IGEs therefore are *expected* to alter the evolutionary potential of a wide range of traits, but whether they do is still unclear. In the wild, trait evolution frequently does not match expectations based on the ratio of direct genetic variance to phenotypic variance and the strength of selection observed in wild populations (Merilä et al., 2001; Pujol et al., 2018). Under artificial selection, the observed response may also not match the expectations based on calculations including only DGEs and the strength of imposed selection (Ellen et al., 2014; Falconer, 1981; Muir, 2005). Studying the role of IGEs in trait evolution more broadly is therefore essential to increase the predictability of evolutionary change and explain these apparent patterns of “evolutionary stasis”, both in the wild and under artificial conditions.

The recent increase in the number of studies providing estimates of IGEs across taxa presents us a timely opportunity to quantitatively assess the role that IGEs play in trait evolution and their importance relative to DGEs. Further, by comparing standardised estimates across contexts and among different trait types, we can assess when IGEs are particularly prevalent, and when trait evolution will diverge most from that predicted by DGEs alone. For example, behavioural traits are a key element of any social interaction and are typically considered to be highly plastic. Behavioural traits may therefore show large IGE variance (Bailey et al., 2018; see Hunt et al., 2019 for a review of IGEs on behaviours). However, behavioural traits may also vary more in response to the non-social environment, in which case the proportion of variance in phenotypes due to IGEs may not be so high. IGEs could also impact traits related to competition such as growth, resource-dependent life-history traits, or fitness components that are not social traits *per se* (Fisher & McAdam, 2019; Wilson, 2014), but still depend on the phenotypes of conspecifics. Many studies on IGEs have taken place on captive populations housed at relatively high densities where individuals cannot avoid social interactions (reviewed in Ellen et al., 2014). In contrast, organisms in the wild typically live at lower densities and may have the option of avoiding social interactions when beneficial. There is therefore the untested possibility that IGEs in the wild are weaker than in captive populations, and so may not contribute as much to the trait’s evolutionary potential, and ultimately, change.

We performed a systematic review and meta-analysis to quantitatively assess the overall magnitude and generality, and several patterns of variation, in IGEs across the published literature. We did not however consider maternal effects, as they have recently been reviewed elsewhere (see Moore et al., 2019). Further, we only analysed effect sizes stemming from the “variance-partitioning” approach used to estimate IGEs (see Methods for further details) since Bailey and Desjonquères (2022) recently reviewed IGEs estimates using the “trait-based” approach. A glossary with definitions of terms used throughout is available in Table 1.

**Table 1.** Glossary of notation used in the manuscript with associated names and formulae.

| Notation | Name | Formula |
| --- | --- | --- |
| DGEs | Direct genetic effects | - |
| IGEs | Indirect genetic effects | - |
| $V_A$ | Direct additive genetic variance | - |
| $V_{IGE}$ | Indirect genetic effects variance | - |
| $V_P$ | Total phenotypic variance | - |
| $h^2$ | Heritability (narrow sense) | $V_A / V_P$ |
| *social* $h^2$ | Social heritability (narrow sense) | $V_{IGE} / V_P$ |
| $I_{DGE}$ | Evolvability | $V_A / \bar{X}^2$ |
| $I_{IGE}$ | Social evolvability | $V_{IGE} / \bar{X}^2$ |
| $COV_{DGE-IGE}$ | DGE-IGE covariance | - |
| $r_{DGE-IGE}$ | DGE-IGE correlation | $COV_{DGE-IGE} / \sqrt{V_A * V_{IGE}}$ |
| $\tau^2$ | Total heritable variance | $[V_A + Cov_{A-IGE} * 2 * (n-1) + V_{IGE} * (n-1)^2] / V_P$ |
| - | “Covariance component” of $\tau^2$ | $(Cov_{A-IGE} * 2 * [n-1])$ |
| - | “IGE variance component” of $\tau^2$ | $(V_{IGE} * [n-1]^2)$ |
| - | Response to individual selection among unrelated individuals | $1 * [V_A + (n-1) Cov_{A-IGE}]$ |
| $n$ | Group size | - |

Our aims were as follows:

1. Quantify the relative magnitude of IGEs across studies (*social h*^2^ = variance in IGEs [V_IGE_] / total phenotypic variance [V_P_]). We had no clear prediction about its magnitude, but based on our understanding of the literature we expected social *h*^2^ to be relatively low.
2. Test whether IGEs differ among (a) types of traits, (b) ages, (c) sexes, and (d) wild and captive populations. Our predictions for (a) were that, because traits closely linked to fitness such as life-history traits typically have low heritabilities, but high evolvabilities (see Hansen et al., 2011; Houle, 1992; Postma 2014, for detailed metric interpretations), the pattern for IGEs variance would be similar. Instead, we expected that behavioural traits would be highly plastic and therefore may show larger IGEs, but also high environmental variances, giving low heritabilities, but high evolvabilities (Bailey et al., 2018). We also predicted for (d) higher effects in captive populations compared to wild populations given the difficulty of avoiding interactions in captivity. We had no specific predictions for the remaining sub-aims.
3. Quantify the magnitude of IGEs in comparison to DGEs. Specifically, we compared: (a) direct additive genetic variance (*V_A_*) vs. *V_IGE_*, (b) heritability (*h*^2^) vs. *social h* ^2^, and (c) *I_DGE_* vs. *I_IGE_*(where *I_DGE_* = *V_A_*/*x̄*^2^ and *I_IGE_* = *V_IGE_*/*x*^2^*̄*; Bailey et al., 2018). The latter relates variances to the trait mean to assess evolvability (i.e. a trait’s evolutionary potential in response to selection, Hansen et al., 2011; Houle, 1992). IGEs are mediated by social interactions, giving a less direct relationship with trait expression than DGEs, and thus we expected them to be smaller than DGEs.
4. Test whether IGEs alter evolutionary trajectories, which we evaluated in two ways: (a) by assessing the overall size and direction of the DGE-IGE correlation (*r*_DGE-IGE_) to understand whether IGEs are expected to increase (positive correlations) or decrease (negative correlations) the rate of evolutionary change in response to direct individual level selection, and (b) by comparing narrow-sense heritability (*h^2^)* with total heritable variance (τ^2^). Since *h^2^* only considers DGEs, while τ^2^ considers DGEs, IGEs, and their covariance, larger τ^2^ compared to h^2^ values would indicate that IGEs increase the amount of genetic variation available to selection and can potentially speed up evolution, while smaller τ^2^ values would indicate IGEs can potentially slow down evolution.

## Materials and methods

### Systematic review

We performed our systematic literature search on the 8^th^ of August 2019 in both *Web of Science Core Collection* (databases: SCI-EXPANDED 1900-present, SSCI 1956-present, A&HCI 1975-present, ESCI 2015-present) and *Scopus*, and consisted of three different steps. First, we searched for studies on IGEs published across all years using the string: ((“indirect genetic effect*”) OR (“social genetic effect*”) OR (“associative genetic effect*”) OR (“interacting phenotype*”)). Second, we performed a backward search to identify references cited in two key recent reviews on the topic: Ellen et al. (2014) and Bailey et al. (2018). Third, we performed a forward search to identify references citing at least one of three seminal theoretical papers on the topic: Bijma & Wade (2008), Moore et al. (1997), and Wolf et al. (1998). We downloaded all the references and found and removed duplicates using the R package ‘revtools’ v.0.3.0 (Westgate, 2019). We screened the titles and abstracts of 1058 unique references using Rayyan (Ouzzani et al., 2016), of which 199 references were selected for full-text screening. Full-text screening identified 111 studies, of which 63 applied an ‘animal model’ approach to analyse their data and were then subjected to data extraction for our meta-analysis. All authors equally performed both the title-and-abstract and the full-text screening using predefined decision trees detailing the inclusion and exclusion criteria (Figures S1 and S2). To reduce any potential observer biases, the title and abstract of 27% of the references (N = 284), and the full text of 22% of the references (N = 44) were screened by more than one observer. Conflicting decisions (title-and-abstract: 7% [N = 20]; full-text: 16% [N = 7]) were collectively discussed and resolved. For more detailed information see our PRISMA diagram (Figure S3; Moher et al., 2009) and the provided data (https://github.com/ASanchez-Tojar/meta-analysis_IGEs).

### Data extraction

Data were extracted from the text and tables in the main text and supplementary materials of the studies identified by our search. All authors equally performed data extraction and the data extraction of all the papers was double-checked by another author. The original and second observer discussed and resolved unclear decisions to ensure reliability. To ensure this process was uniform, all authors extracted data from four selected papers as a pilot at the beginning of this step.

From each included study, we extracted information about the article (year of publication, journal, author information), study subjects (species, population and years studied, population type [captive, semi-captive {defined as a population kept in captivity for no more than five generations}, wild], sex [males, females, both] and age [juveniles, adults, both] of included subjects) and type of trait(s) studied (behaviour, development, metabolism and physiology, morphology, reproduction, survival). In addition, we extracted the information needed to interpret the statistical estimates of each animal model, namely whether response variables were mean-or variance-standardized, the error distribution and any transformations used, mean group size used to estimate IGEs, mean and standard deviation (SD) of the trait(s) studied (i.e. the response variables), and whether fixed effects pertaining to focal or partner individual traits were included in the models. Lastly, we extracted sample sizes (e.g. number of individuals studied, number of observations), pedigree-related information, and the estimates from the animal models (e.g. direct [*V_A_*] and indirect genetic effect variances [*V_IGE_*], narrow-sense [*h*^2^] and total heritability [τ^2^], direct-indirect genetic correlation [*r* _IGE_], and direct-indirect genetic covariance [*CoV*_A-IGE_]).

After data extraction, we excluded 16 studies: (i) 8 studies (k = 13 effect sizes) that did not contain usable data (i.e. because of being a conference abstract, reporting the same data as other studies, not estimating IGEs using the animal model approach, or solely reporting estimates from existing literature), (ii) 1 study (k = 1 effect size) on humans, (iii) 2 studies (k = 7 effect sizes) on *Eucalyptus globulus* because we did not find data for any other plant species, and (iv) 5 studies (k = 10 effect sizes) that exclusively analysed non-Gaussian traits. Notably, several wild studies fell into this latter exclusion category, reducing the sample size for that category (see below). Requests for missing or partially reported data were sent to the corresponding authors of 38 studies using a standardized email template (provided in Supplementary Materials), from which we obtained additional data for 25 studies. Finally, note that our meta-analytic dataset included 13 out of the 21 studies included in the qualitative literature review by Ellen et al. (2014; see Table 4 therein), with the difference being explained by the inclusion/exclusion criteria explained above. For a complete list of studies and extracted variables see Table 2 and the provided data (https://github.com/ASanchez-Tojar/meta-analysis_IGEs).

**Table 2.** List of studies (n = 47) from which data was extracted and included in our meta-analysis on indirect genetic effects.

| Study ID | Year | Journal | Number of effect sizes (k) | Species | Reference |
| --- | --- | --- | --- | --- | --- |
| IGE0057 | 2015 | Genetics Selection Evolution | 1 | *Penaeus vannamei* | (Luan et al., 2015) |
| IGE0144 | 2013 | Agriculturae Conspectus Scientificus | 3 | *Sus scrofa* | (Rostellato et al., 2013) |
| IGE0195 | 2012 | Journal Of Animal Science | 3 | *Sus scrofa* | (Duijvesteijn et al., 2012)* |
| IGE0196 | 2016 | Journal Of Animal Ecology | 1 | *Melospiza melodia* | (Germain et al., 2016) |
| IGE0197 | 2016 | Evolution | 1 | *Tribolium castaneum* | (Ellen et al., 2016) |
| IGE0199 | 2019 | Journal Of Evolutionary Biology | 3 | *Chorthippus biguttulus* | (Chakrabarty et al., 2019) |
| IGE0202 | 2015 | Journal Of Animal Science | 10 | *Sus scrofa* | (Rostellato et al., 2015) |
| IGE0203 | 2014 | Genetics Selection Evolution | 19 | *Gadus morhua* | (Nielsen et al., 2014)* |
| IGE0242 | 2007 | Journal Of Animal Science | 8 | *Bos taurus* | (Van Vleck et al., 2007)* |
| IGE0243 | 2010 | Journal Of Animal Science | 3 | *Sus scrofa* | (Hsu et al., 2010)* |
| IGE0253 | 2012 | Journal Of Animal Science | 1 | *Bos taurus* | (Sartori & Mantovani, 2012) |
| IGE0291 | 2008 | Journal Of Animal Science | 1 | *Sus scrofa* | (Chen et al., 2008)* |
| IGE0292 | 2016 | Journal Of Animal Breeding And Genetics | 3 | *Neovison vison* | (Alemu et al., 2016) |
| IGE0294 | 2005 | Journal Of Animal Science | 1 | *Sus scrofa* | (Arango et al., 2005) |
| IGE0347 | 1997 | British Poultry Science | 6 | *Gallus gallus* | (Kjaer & Sørensen, 1997) |
| IGE0380 | 2016 | Aquaculture | 1 | *Oreochromis niloticus* | (Khaw et al., 2016) |
| IGE0385 | 2019 | Asian-Australasian Journal Of Animal Sciences | 1 | *Sus scrofa* | (Hong et al., 2019) |
| IGE0391 | 2013 | Journal Of Animal Science | 3 | *Sus scrofa* | (Bergsma et al., 2013)* |
| IGE0406 | 2005 | Proceedings Of The National Academy Of Sciences Of The United States Of America | 16 | *Drosophila serrata* | (Petfield et al., 2005) |
| IGE0468 | 2019 | Behavior Genetics | 24 | *Mus musculus* | (Dewan et al., 2019) |
| IGE0483 | 2018 | Evolution | 6 | *Gryllus bimaculatus* | (Han et al., 2018) |
| IGE0493 | 2019 | Ecology Letters | 1 | *Tamiasciurus hudsonicus* | (Fisher, Haines, et al., 2019) |
| IGE0494 | 2019 | Animal | 1 | *Sus scrofa* | (Ragab et al., 2019) |
| IGE0501 | 2009 | Proceedings Of The Royal Society B-Biological Sciences | 5 | *Peromyscus maniculatus sonoriensis* | (Wilson et al., 2009)* |
| IGE0505 | 2014 | Genetics Selection Evolution | 3 | *Neovison vison* | (Alemu et al., 2014)* |
| IGE0507 | 2012 | Genetics | 4 | *Gallus gallus* | (Peeters et al., 2012)* |
| IGE0509 | 2010 | Journal Of Evolutionary Biology | 1 | *Chroicocephalus scopulinus* | (Teplitsky et al., 2010) |
| IGE0515 | 2017 | Scientific Reports | 1 | *Gryllus bimaculatus* | (Santostefano et al., 2017) |
| IGE0537 | 2017 | Genetics Selection Evolution | 2 | *Oryctolagus cuniculus* | (Piles et al., 2017) |
| IGE0572 | 2018 | Genetics Selection Evolution | 5 | *Oryctolagus cuniculus* | (David et al., 2018) |
| IGE0598 | 2010 | Journal Of Animal Science | 1 | *Sus scrofa* | (Bouwman et al., 2010)* |
| IGE0640 | 2014 | Poultry Science | 2 | *Gallus gallus* | (Sun et al., 2014) |
| IGE0654 | 2007 | Genetics | 1 | *Gallus gallus* | (Bijma et al., 2007) |
| IGE0657 | 2013 | Evolution | 1 | *Coturnix japonica* | (Muir et al., 2013)* |
| IGE0746 | 2014 | Genetics Selection Evolution | 8 | *Gallus gallus* | (Brinker et al., 2014)* |
| IGE0752 | 2015 | Genetics Selection Evolution | 4 | *Gallus gallus* | (Brinker et al., 2015) |
| IGE0786 | 2017 | Animal Reproduction | 1 | *Sus scrofa* | (Hong et al., 2017) |
| IGE0824 | 2014 | Evolution | 2 | *Drosophila melanogaster* | (Edward et al., 2014) |
| IGE0852 | 2019 | Journal Of Evolutionary Biology | 1 | *Tamiasciurus hudsonicus* | (Fisher, Wilson, et al., 2019) |
| IGE0857 | 2015 | Proceedings Of The Royal Society B: Biological Sciences | 1 | *Aegithalos caudatus* | (Adams et al., 2015) |
| IGE0858 | 2018 | Genetics Selection Evolution | 2 | *Sus scrofa* | (Nielsen et al., 2018) |
| IGE0889 | 2008 | Poultry Science | 3 | *Gallus gallus* | (Ellen et al., 2008)* |
| IGE0920 | 2008 | Genetics | 5 | *Sus scrofa* | (Bergsma et al., 2008) |
| IGE0927 | 2017 | Heredity | 1 | *Sus scrofa* | (Canario et al., 2017) |
| IGE1023 | 2013 | Genetics Selection Evolution | 2 | *Gallus gallus* | (Peeters et al., 2013) |
| IGE1038 | 2019 | Scientific Reports | 6 | *Sus scrofa* | (Wu et al., 2019) |
| IGE1050 | 2008 | Evolution | 1 | *Larus canus* | (Brommer & Rattiste, 2008) |

\* References included in the qualitative review performed by Ellen et al. (2014: Table 4 in that review).

### Effect size calculation

Our effect sizes of interest were: (i) relative magnitude of DGEs (narrow-sense heritability or *h^2^*) and IGEs (*social h^2^*; aims 1, 3 and 4), (ii) unstandardized variances associated with DGEs (*V_A_*) and IGEs (*V_IGE_*), which we log-transformed prior to the analyses to better match model assumptions (aim 3), (iii) DGE and IGE evolvability (*I_DGE_* and *I_IGE_*; aim 3), (iv) DGE-IGE correlation (*r*_DGE-IGE_; aim 4), which when not reported, was calculated as *COV*_A-IGE_ divided by the square root of the product of *V_A_* and *V_IGE_*, and (v) total heritable variance, τ^2^ (i.e. the ratio of total heritable variation and phenotypic variance, aim 4), which was calculated as [V + Cov *2*(n-1) + V *(n-1)^2^] / V_P_, where n is group size (Bijma, 2010; Bijma et al., 2007). As in previous meta-analyses (e.g. Dochtermann et al. 2019, Holtmann et al. 2017), for most of our analyses we treated intra-class correlations as equivalent to Pearson’s correlation coefficients (*r*) and applied Fisher’s *Z* transformation to obtain our main effect size of interest (i.e. *Zr* and its associated sampling variance, *VZr*). Therefore, those meta-analytic estimates correspond to *Zr* instead of *r* throughout; however, we present back-transformed values (i.e. to *r*) in the figures to aid biological interpretation. The variance in *Zr* (i.e. sampling variance) was calculated as: *VZr* = 1 / (n-3), where n represents the number of unique individuals phenotyped. The different levels of replication analyzed (e.g., number of groups, number of replicates per group) in the original mixed-effects models from which we obtained the estimates may lead to an underestimation of sampling variance. To test the robustness of our results to an increase in sampling variance, we performed sensitivity analyses by rerunning several analyses after having artificially increased sampling variance by a factor of 10 (i.e., *VZr*_new_ = *VZr*_original_ * 10). This increase did not substantially change our conclusions (Supplementary Material). If *social h ^2^* was not directly reported in the corresponding study, we calculated it ourselves as V_IGE_ / V_P_. If V_P_ was not reported, we calculated it ourselves as *V_A_* / *h*^2^ when these variables were reported, and then calculated *social h^2^*. These calculations allowed us to add 13 additional effect sizes to our dataset. Additionally, there were nine estimates of *V_IGE_* reported as “0” in the primary literature. To avoid removing those values (as zeros cannot be log-transformed), for the comparison of *V_A_* and *V_IGE_* only (aim 3) we converted the reported *V_IGE_* values to the minimum possible rounded value based on how the “0” was reported (e.g. reported 0.00 transformed to 0.001). We note that small variances in our dataset might be upwardly biased due to the likely prevalence of estimates from REML models and/or the likely preference for reporting means and medians rather than modes in Bayesian models (see Pick et al. 2023).

### Meta-analyses and meta-regressions

We first ran an intercept-only meta-analytic model to estimate the overall effect size (i.e. meta-analytic mean) for *social h*^2^. We then ran a meta-regression for each of our hypotheses, giving eight meta-regressions, each containing a single moderator at a time. All meta-regressions were complete-case meta-regressions (i.e. effect sizes with unreported moderator data were discarded). In addition, we also ran an intercept-only meta-analysis to estimate the overall effect size for *r*_DGE-IGE_.

All models included the following six random effects: (i) phylogeny, which consisted of a phylogenetic relatedness correlation matrix (Cinar et al., 2022), (ii) species ID, encompassing effect sizes obtained from the same species and accounts for among-species differences not explained by phylogeny, (iii) population ID, encompassing effect sizes obtained from the same population, (iv) study ID, encompassing effect sizes extracted from the same study, (v) group ID, encompassing effect sizes derived from the same group of animals and allowing us to model different experimental groups within the same study, and (vi) record ID (unit-level effect), which represents residual and/or within-study variance in meta-analytic models. The phylogeny was built by searching for the species in the Open Tree Taxonomy (Rees & Cranston, 2017) and retrieving their phylogenetic relationships from the Open Tree of Life (Hinchliff et al., 2015) using the R package ‘rotl’ v.3.0.11 (Michonneau et al., 2016). We estimated branch lengths following (Grafen, 1989) as implemented in the function ‘compute.brlen()’ of the R package ‘ape’ v.5.4-1 (Paradis & Schliep, 2019) and included the phylogenetic variance–covariance matrix as a random effect in all models. For the meta-regressions used to compare the (relative) magnitude of DGEs and IGEs as well as to compare narrow-sense heritability (*h^2^*) to total heritable variance (τ^2^; hypotheses 3a, 3b, 3c and 4b), since each pair of estimates (e.g. V_IGE_ and V_A_) was obtained from the same model, we included an additional random effect (vii; model ID) that encompassed effect sizes obtained from the same animal model to account for that extra level of nonindependence.

To evaluate the importance of each potential source of variation, for all meta-analytic (intercept-only) models, we calculated four heterogeneity metrics (*σ^2^*, *I^2^*, *CV*, *M*) to understand total and random effect-specific heterogeneities following the pluralistic approach suggested by Yang et al. (2023). For all meta-regressions, we estimated the percentage of heterogeneity explained by the moderator as *R*^2^ (Nakagawa & Schielzeth, 2013).

We used the programming language R v.4.0.4 (R Core Team, 2021) throughout. All meta-analytic and meta-regression models were fitted in the R package ‘metafor’ v.2.5-82 (Viechtbauer, 2010). Results were graphically represented as orchard plots using the R package ‘orchaRd’ v.2.0 (Nakagawa et al., 2023).

### Testing for sources of bias, including publication bias

We confirmed the robustness of our results by running several sensitivity analyses that tested: a) the effect of including fixed effects pertaining to social partners in the original animal models, b) whether conclusions remained when only using originally reported estimates (i.e. excluding our own calculations of some effect sizes; see above), c) evidence for publication bias both small-study and decline effects (following Nakagawa et al., 2022; Sánchez-Tójar et al., 2018) and d) the effect of potentially underestimating sampling variance. None of these factors proved to be significant sources of bias, see the Supplementary Materials for further details on our approach and the results.

## Results

Our full dataset contained 180 effect sizes from 47 studies published between 1997 and 2019 and involving 21 species. The median and mean number of studies per species was 1 and 2.2, respectively, with *Sus scrofa* (N = 15 studies) and *Gallus gallus* (N = 8 studies) being the most commonly studied species. Note that since we performed complete-case analyses throughout, sample sizes differ slightly across models and research questions.

### Aim 1 What is the relative magnitude of IGEs (social h2)?

The intercept-only model revealed a small, but statistically significant estimate for social *h*^2^ (*Zr* = 0.031, 95% confidence intervals, CI, [0.012, 0.051], 95% prediction intervals, PI, [-0.061, 0.124], *p* = 0.002; k = 146 effect sizes, N = 40 studies, 19 species; Figure 2), but heterogeneity was high (*I^2^_total_* = 93.2%; *CV_total_*= 1.46; *M_total_* = 0.70). *I^2^_total_* and *CV_total_* indicated that heterogeneity in our dataset is, on average, around 14 times larger than statistical noise and around 1.5 times larger than the meta-analytic mean. Random effect-specific *I^2^* estimates showed that 62.9% of heterogeneity was attributed to study ID, 20.1% to species ID not linked to the phylogenetic tree, and 10.2% to record ID. Negligible or zero relative heterogeneity was attributable to the phylogenetic tree, group ID, and population ID.

**Figure 2.**
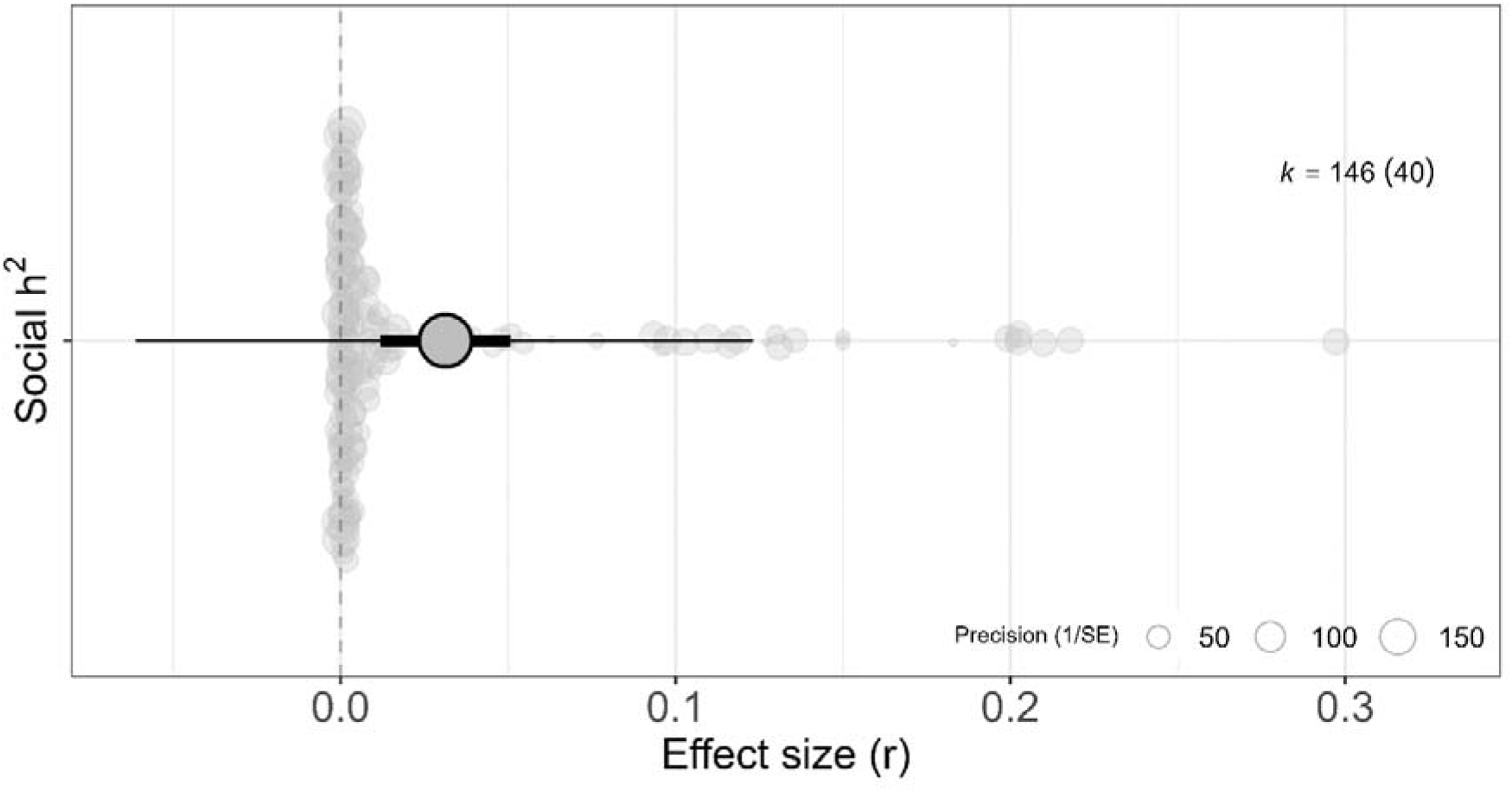
Social heritability (*social h*^2^) from a phylogenetic multilevel meta-analysis with 40 studies and 146 effect sizes (Aim 1). Orchard plot showing the back-transformed meta-analytic mean as *r*, 95% confidence intervals (thick whisker), 95% prediction intervals (thin whisker) and individual effect sizes (*r*) scaled by their precision (semitransparent circles). *k* corresponds to the number of effect sizes, the number of studies is shown in brackets. Note that uncertainty measures (i.e., 95% CI and 95% PI) are generated assuming a normal distribution around the meta-analytic mean, explaining why some intervals may overlap zero substantially despite the data analyzed being bounded between 0 and 1.

### Aim 2 Does the magnitude of IGEs vary with intrinsic or extrinsic factors?

#### 2a Trait category

Social *h*^2^ differed among trait categories (Figure 3a) with the moderator explaining 32.1% of the heterogeneity (*R^2^_marginal_* = 0.321). The only trait categories with a mean estimate statistically significantly different from zero were behaviour (*Zr* = 0.061, 95% CI [0.032, 0.091]) and reproduction (*Zr* = 0.053, 95% CI [0.014, 0.093]; Figure 3a). Behavioural traits had statistically significantly higher social *h^2^* values than all other trait categories except for reproduction and survival (see Table S1 for p-values from all pair-wise tests). The post-hoc Wald tests showed that most of the differences between the remaining trait categories were not statistically significant, with only reproduction either being significantly higher than both development and morphology, or showing a trend in that direction when compared to metabolism and physiology (Table S1).

**Figure 3.**
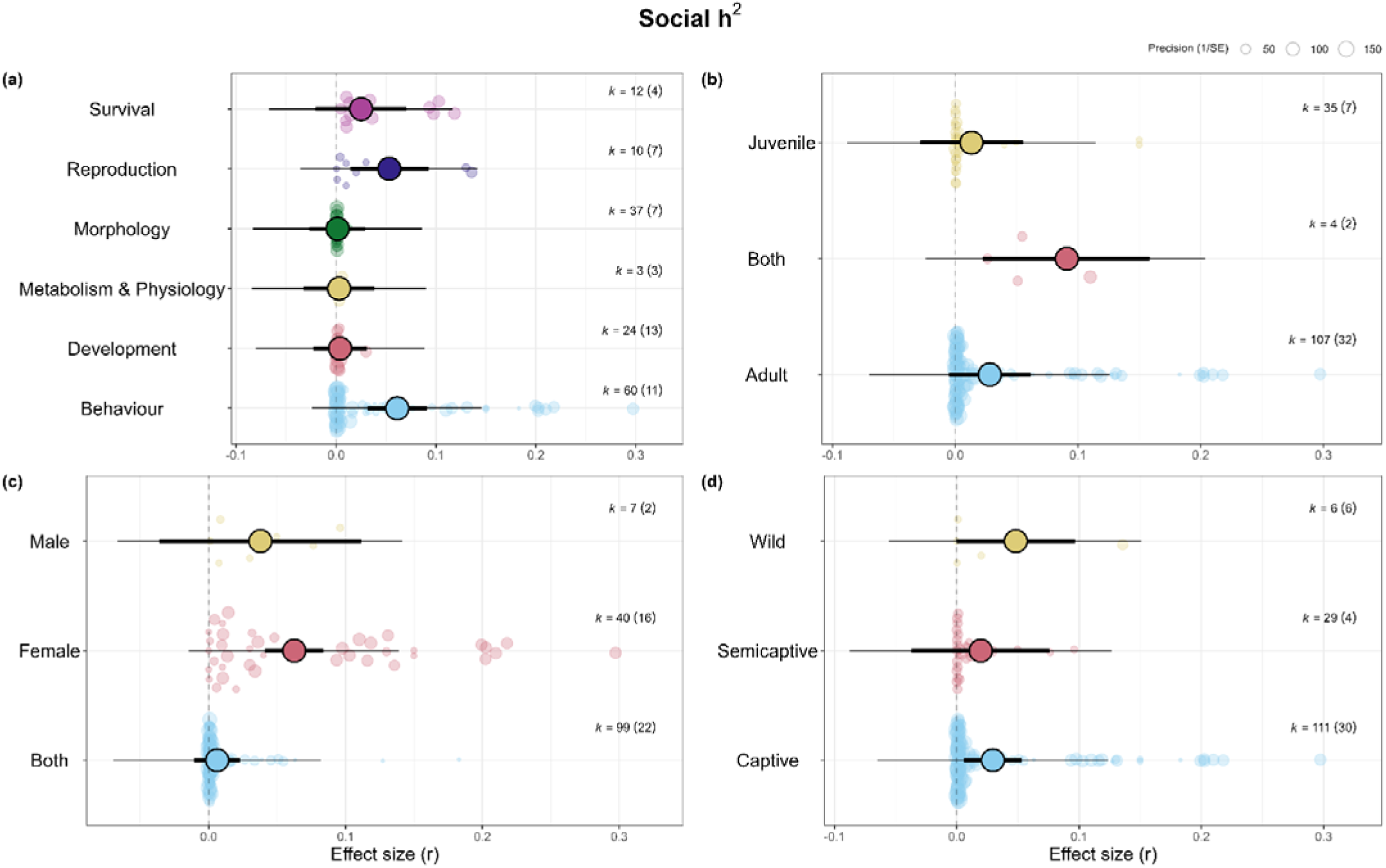
Social heritability (*social h* ^2^) meta-analytic estimates from phylogenetic multilevel uni-moderator meta-regressions testing the following moderators (Aim 2): (a) Trait category, (b) Age, (c) Sex, (d) Population type. Orchard plot showing the back-transformed meta-analytic mean as *r*, 95% confidence intervals (thick whisker), 95% prediction intervals (thin whisker) and individual effect sizes (*r*) scaled by their precision (semitransparent circles). *k* corresponds to the number of effect sizes, the number of studies is shown in brackets. Note that uncertainty measures (i.e., 95% CI and 95% PI) are generated assuming a normal distribution around the meta-analytic mean, explaining why some intervals may overlap zero substantially despite the data analyzed being bounded between 0 and 1.

#### 2b Age

Social *h*^2^ differed among age categories and was statistically significantly larger for effect sizes combining both age classes (*Zr* = 0.091, 95% CI [0.022, 0.160]) compared to those from juveniles-only (*Zr* = 0.013, 95% CI [-0.029, 0.055], p = 0.049) and adults-only (*Zr* = 0.028, 95% CI [-0.05, 0.062], p = 0.026; Table S1, Figure 3b). Estimates for juveniles-only and adults-only were not statistically significantly different from each other (p = 0.403; Table S1, Figure 3b). Overall, the moderator age explained 6.8% of the heterogeneity (*R^2^_marginal_* = 0.068). Note that there were only four effect sizes for the category combining both age classes, and therefore caution should be applied when drawing conclusions from those differences.

#### 2c Sex

Social *h*^2^ differed among sex categories, with social *h*^2^ for females-only effect sizes being larger (*Zr* = 0.063, 95% CI [0.041, 0.084]) than when including both sexes (*Zr* = 0.006, 95% CI [-0.011, 0.023], p < 0.001) or for only males (though not statistically significantly so; *Zr* = 0.038, 95% CI [-0.036, 0.112], p = 0.524; Table S1, Figure 3c). Sex as a moderator explained a large amount of heterogeneity (31.1%; *R^2^_marginal_* = 0.311); however, there were only seven effect sizes for male-only data, and, out of the 40 female-only effect sizes, 28 were from chickens in a production setting, which limits our ability to generalise these results.

#### 2d Population type

Social *h*^2^ did not differ among population types (Table S1, Figure 3d) with the moderator explaining 1.5% of the heterogeneity (*R^2^_marginal_* = 0.015). Social *h*^2^ was statistically significantly different from zero in captive populations (*Zr* = 0.030, 95% CI [0.006, 0.053]), however, the estimate did not statistically differ from zero in semicaptive populations (*Zr* = 0.020, 95% CI [−0.037, 0.076]) and only marginally so for wild populations (*Zr* = 0.048, 95% CI [−0.000, 0.097]: Table S1, Figure 3d). The pair-wise comparisons revealed no statistically significant differences between all three levels (*p* > 0.448 in all cases; Table S1, Figure 3d). Note, however, that there were only six studies (k = 6 effect sizes) from wild populations and four (k = 29 effect sizes) from semicaptive populations.

### Aim 3 What is the relative magnitude of IGEs compared to DGEs?

We ran three meta-regressions to test for the difference in magnitude between V_IGE_ and V_A_ (aim 3a); social *h*^2^ and *h*^2^ (aim 3b); and *I* and *I* (aim 3c). We found that regardless of whether IGEs were estimated as raw variance (log(V_IGE_) = -3.775, 95% CI [-6.562, -0.988] vs. log(V_A_) = -0.019, 95% CI [-2.806, 2.767], p < 0.001), variance-standardized variance (*social h^2^*: *Zr* = 0.042, 95% CI [-0.027, 0.111] vs. *h*^2^: *Zr* = 0.263, 95% CI [0.194, 0.331], p < 0.001), or mean-standardized variance (*I*_IGE_ = 0.045, 95% CI [-0.134, 0.223] vs. *I*_DGE_ = 0.224, 95% CI [0.045, 0.402], p = 0.003), IGEs were smaller than DGEs (Figure 4). The moderator explained a considerable amount of variance in each meta-regression (13.5%, 26.8%, and 4.0%, respectively).

**Figure 4.**
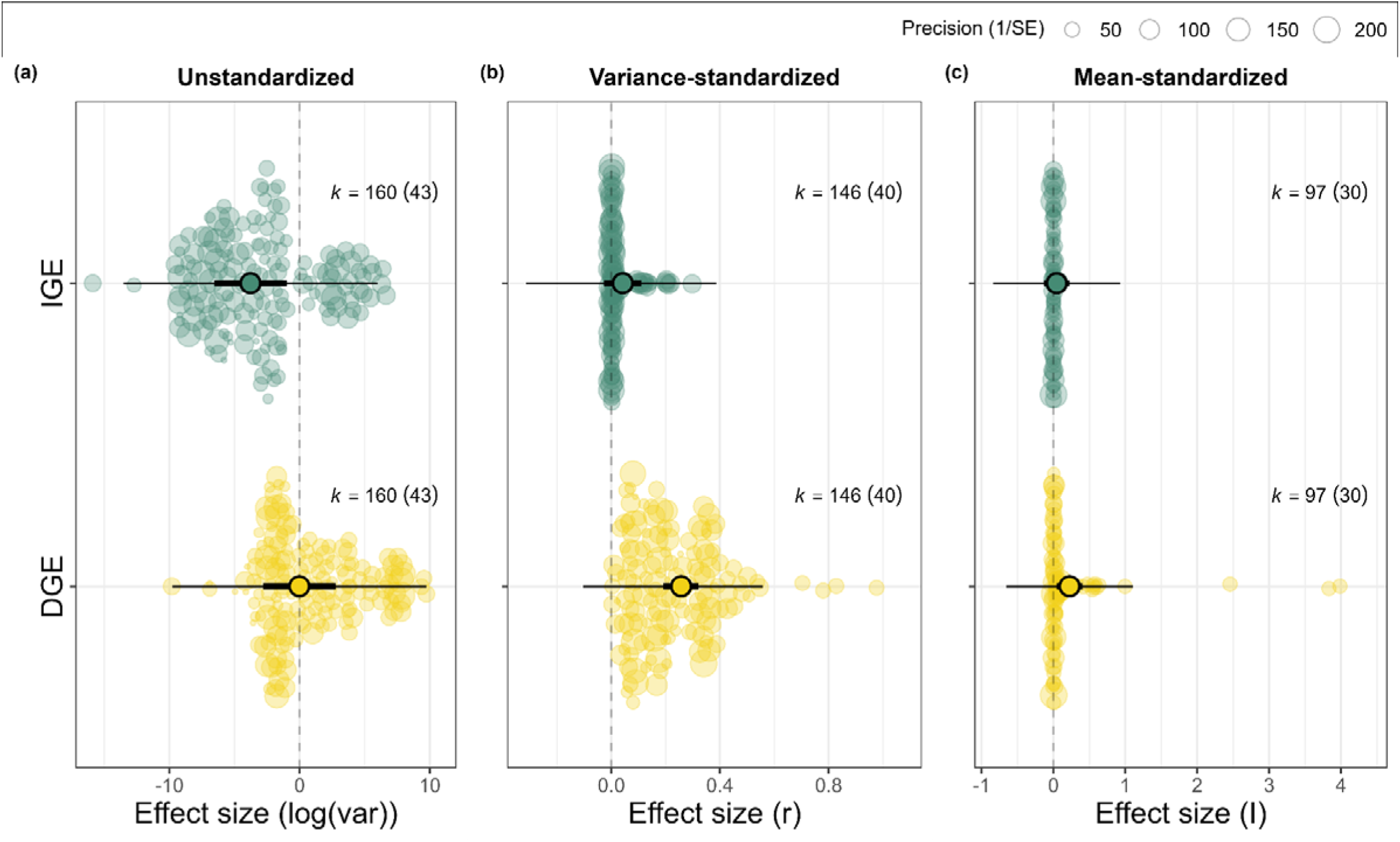
IGEs and DGEs meta-analytic estimates from phylogenetic multilevel uni-moderator meta-regressions (Aim 3) comparing: (a) *V*_A_ vs. *V*_IGE_, (b) narrow-sense *h*^2^ vs. *social h*^2^, and (c) *I* vs. *I* (evolvability). Orchard plot showing the back-transformed meta-analytic mean as *r*, 95% confidence intervals (thick whisker), 95% prediction intervals (thin whisker) and individual effect sizes (*r*) scaled by their precision (semitransparent circles). *k* corresponds to the number of effect sizes, the number of studies is shown in brackets. Note that for (b) and (c) uncertainty measures (i.e., 95% CI and 95% PI) are generated assuming a normal distribution around the meta-analytic mean, explaining why some intervals may overlap zero substantially despite the data analyzed being bounded between 0 and 1.

### Aim 4 Do IGEs alter evolutionary trajectories?

The intercept-only model revealed an overall positive but statistically non-significant correlation between DGEs and IGEs (*r*_DGE-IGE_; Z*r* = 0.270, 95% CI [-0.081, 0.621]; Figure 5b), but heterogeneity was high (*I^2^_total_*= 99.98; *CV_total_* = 3.25; *M_total_* = 0.85). This high heterogeneity explains the large 95% prediction intervals observed (95% PI [-1.497, 2.036]), which shows that, although positive on average, *r*_DGE-IGE_ should be expected to range from very negative to very positive depending on the particular context investigated.

**Figure 5.**
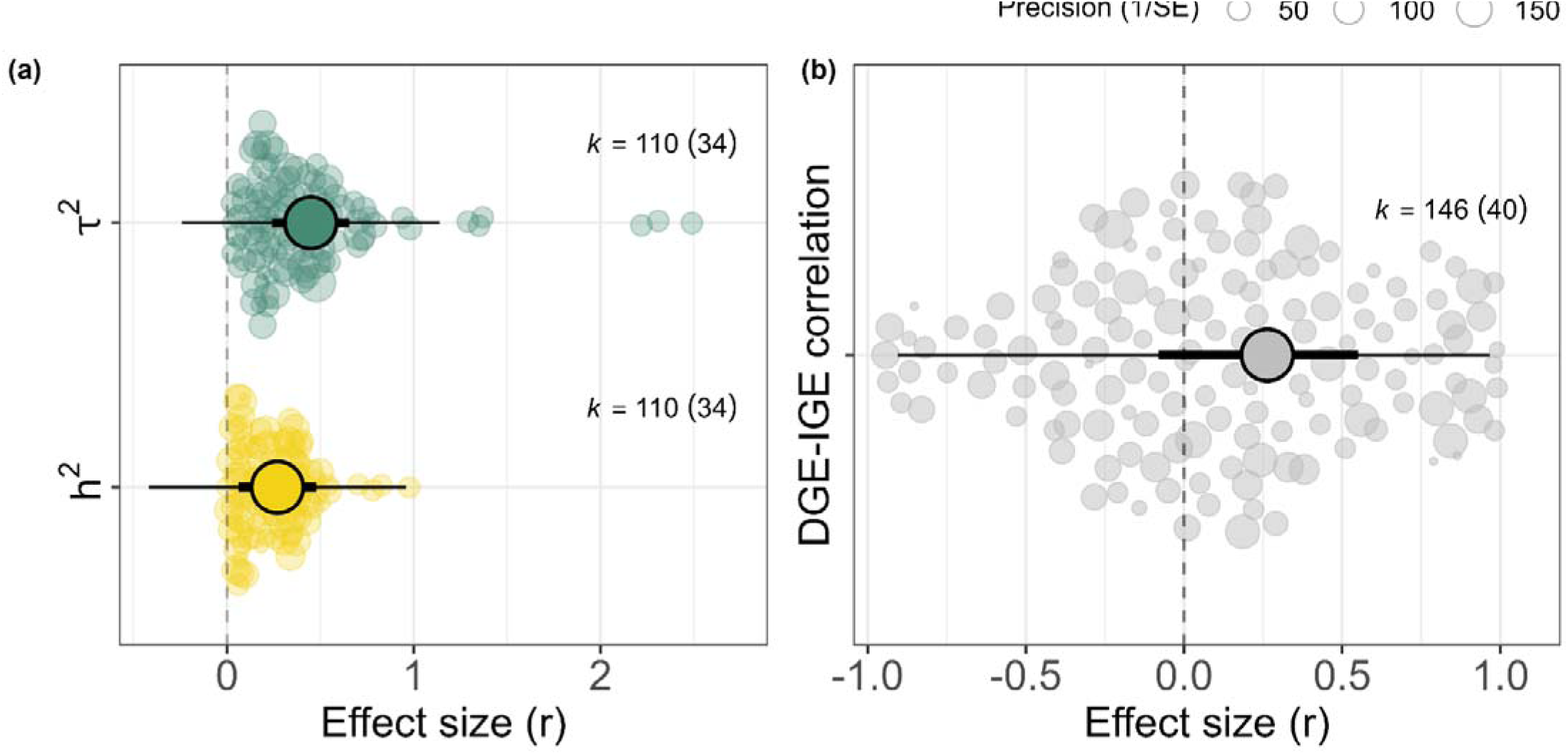
Meta-analytic estimates for Aim 4: (a) phylogenetic multilevel meta-regression comparing narrow-sense heritability (*h*^2^) to total heritable variance (τ); (b) phylogenetic multilevel meta-analysis of *r*_DGE-IGE_. Orchard plot showing the original (a) and back-transformed meta-analytic mean as *r*, 95% confidence intervals (thick whisker), 95% prediction intervals (thin whisker) and individual effect sizes (*r*) scaled by their precision (semitransparent circles). *k* corresponds to the number of effect sizes, the number of studies is shown in brackets. Note that for (a) uncertainty measures (i.e., 95% CI and 95% PI) are generated assuming a normal distribution around the meta-analytic mean, explaining why some intervals may overlap zero substantially despite the data analyzed being bounded between 0 and 1.

Total heritable variance (τ^2^) was stastistically significantly larger (*r* = 0.449, 95% CI [0.241, 0.656]) than narrow-sense heritability (*h*^2^; *r* = 0.270, 95% CI [0.062, 0.478], p < 0.001) and the moderator explained 6.7% of the heterogeneity (*R^2^_marginal_* = 0.067; Figure 5a).

To better understand whether the increase in τ^2^ compared to *h*^2^ was driven by the IGEs variance and its relationship with group size (*n*) or due to the seemingly positive *r*_DGE-IGE_ detected, as a post-hoc investigation we calculated the two components of τ^2^ that stem from V_IGE_, following Bijma et al., 2007 (*eq. 6*) and qualitatively compared their overall magnitude. These are the “covariance component” (Cov_A-IGE_*2*[*n*-1]) and the “IGE variance component” (V_IGE_*[*n*-1]^2^). This additional analysis showed that the IGEs variance component was seemingly larger (mean = 693.14) than the covariance component (mean = 241.75), suggesting that it is the IGEs variance component that drives the increase in τ^2^.

In addition, τ^2^ represents how much genetic variation is available to *all* types of selection (including group-level selection). To test how much difference IGEs make to potential responses to direct, individual-level only selection, we calculated the difference between the predicted response to direct selection on unrelated individuals in the presence and absence of IGEs, following eq. 15 in Bijma and Wade (2008), assuming that individuals interact in groups of size *n* and a direct selection gradient of 1 (i.e. 1* [V_A_ + (*n*-1) Cov_A-IGE_]). This post-hoc analysis showed that the response to direct, individual-level only selection among unrelated individuals was not different in the presence of IGEs (Z*r* = 0.183, 95% CI [0.048, 0.319]) and in their absence (Z*r* = 0.225, 95% CI [0.089, 0.360], p = 0.470, *R^2^_marginal_* = 0.002).

## Discussion

We performed a meta-analysis to assess the overall strength and patterns of variation in indirect genetic effects across the published literature. The magnitude of IGEs is small but statistically significant, with social *h^2^* accounting for, on average, around 3% of the phenotypic variation. A recent meta-analysis concluded that genetic and non-genetic maternal effects together explained 11% of the phenotypic variance (Moore et al., 2019). Maternal *genetic* effects alone (a type of IGEs excluded from our own meta-analysis) must be smaller than 11% and, therefore, likely comparable to our IGEs estimate. The high levels of heterogeneity among effects that we found show that, as expected, the importance of IGEs varies across contexts, with the 95% prediction intervals suggesting that once estimate precision is accounted for, *social h^2^* is expected to range from 0 to 0.12 in most cases. Indeed, we found that factors such as trait type as well as age and sex of individuals explain substantial heterogeneity in *social h^2^* estimates across studies. Notably, IGEs were more prominent for behaviours (0.06) and traits related to reproduction (0.05), supporting the hypothesis that labile traits are most strongly affected by social interactions. Furthermeore, as expected, IGEs were smaller in magnitude compared to DGEs, with estimates of direct narrow-sense heritability (*h^2^* = 0.27) being, on average, 6 to 8 times (depending on the statistical model) larger than *social h^2^*. However, despite this large difference, accounting for IGEs and their correlations with DGEs (i.e. total heritable variance, τ*^2^*_)_ leads to a 66% increase in the total amount of genetic variation available to selection (τ*^2^* = 0.45) when compared to narrow-sense *h^2^*. This difference is mostly driven by the relationship between the contribution of IGEs to τ^2^ and group size, but also due to the overall positive correlation bewtween DGEs and IGEs (*r*_DGE-IGE_ = 0.26; although not statistically different from zero). We detected no evidence of publication bias, and our results were robust to data imputations, potential underestimation of the sampling variance, and several aspects of model structure in the original studies (see Supplementary Materials). Overall, our results show the general importance of IGEs for animal populations, the consequences of which we outline below.

### IGEs can enhance the evolutionary potential of traits

The most impactful consequence of IGEs on traits evolution to emerge from our meta-analysis is that IGEs enhance evolutionary potential by increasing the amount of genetic variation available to selection. In their review, Ellen et al., (2014) suggested that total heritable variance (τ^2^) seems to be larger than narrow-sense heritability (*h^2^*) in most domesticated populations; we here confirm this quantitatively and statistically. Two separate but not mutually exclusive mechanisms appear to facilitate this increase in adaptive potential: (1) the influence of IGEs on many interaction partners, and (2) a positive DGEs-IGEs correlation (*r*_DGE-IGE_).

With a post-hoc analysis, we showed that the increase in τ^2^ compared to *h*^2^ is mainly driven by the IGEs variance component (V_IGE_[*n*-1]^2^) rather than covariance component (Cov *2*[*n*-1]). This result suggests that there may be greater evolutionary potential when organisms interact in larger groups, as then the impact of V_IGE_ is magnified; however, V_IGE_ may also decrease, if the influence of any two random individuals on each other is diluted as n increases (see Arango et al., 2005; Hadfield & Wilson, 2007; Wade et al., 2010). Bijma (2010) presents a general model for the relationship of IGEs with group size, but we could not determine where studies included in our meta-analysis used it (but see Duijvesteijn et al. 2012 and Canario et al. 2017 for examples). The suggested importance of group size supports results from previous modelling approaches revealing that both DGEs and IGEs can depend largely on both group size and interaction strength (Liu & Tang, 2016; Trubenová & Hager, 2012). How interactions occur and whether group size will enhance or dilute IGEs will be specific to the biology of the species and the setup of the study. In most captive studies, experimenters enforce one group size across the experiment. Therefore, caution should be exercised when trying to extrapolate estimates of IGEs to situations where group sizes are outside the range of those used to estimate the IGEs in the first place (see also attempts to weight interaction strengths with, for example, social- or spatial-based neighbour networks, and determine effective group size: Bijma, 2010; Costa E Silva et al., 2013; Fisher et al., 2019; Santostefano et al., 2021). Furthermore, differences among individuals in their effect on group mates (e.g. “keystone” individuals: Modlmeier et al., 2014; or among individual variation in “social impact”: Araya-Ajoy et al., 2020; de Groot et al., 2023) could also become important and modulate the magnitude of the effect of IGEs in a group.

The overall positive (though uncertain) correlation between DGEs and IGEs (r_DGE-IGE_ = 0.26) also indicates that heritable social effects tend to facilitate trait evolution rather than impede it. This suggests that IGEs may not be a general explanation for phenotypic stasis in the face of directional selection (Kruuk et al., 2008; Merilä et al., 2001; Pujol et al., 2018; Wilson, 2014). Our result aligns with a review of the interacting phenotypes coefficient Ψ (Bailey & Desjonquères, 2022), which quantifies the effect of a trait on the expression of the same or a different trait in social partners. The sign of Ψ was generally positive, indicating positive feedback between the trait expression in interacting partners, and suggesting a predicted response to selection that is *greater* than in traits where Ψ is zero or without IGEs (Wolf et al., 1998). We then estimated the potential response to selection as an additional comparison for narrow-sense *h*^2^ and τ^2^, which revealed an important caveat. The response to individual-level direct selection in the presence of IGEs (in the absence of relatedness and multilevel selection) was not statistically different from that predicted in their absence. This lack of statistical difference, in contrast to the results for narrow-sense *h*^2^ and τ^2^, appears because it is non-zero relatedness and multilevel selection that magnify the influence of V_IGE_ on the response to selection. Relatedness between the focal individual and the individuals affecting its fitness is crucial for harnessing the heritable variation for adaptive evolution (see Bijma 2011 for further discussion). Therefore, if relatives interact, the contribution of V_IGE_ to traits evolution will begreater. That IGEs do not enhance the response to selection when only individual-level selection occurs may partially explain the surprising lack of response to artificial selection for improvement of socially-affected traits (Craig & Muir, 1996; Ellen et al., 2014; Wade, 1976, 1977), and suggests that multilevel selection may be necessary to “unlock” the evolutionary potential of IGEs (Bijma & Wade, 2008; Muir et al., 2013). However, multilevel selection is not typically quantified in studies of selection in wild populations, nor always applied in breeding designs. We therefore do not yet know the general prevalence of multilevel selection and how that impacts the evolutionary potential of populations, and call researchers in the field to consider quantifying its influence in future studies.

### IGEs are smaller than DGEs

As expected, DGEs were larger than IGEs (e.g. narrow-sense *h^2^* [0.27] was more than 6 times larger than *social h*^2^ [0.03]) across metrics. Our estimates of *h^2^* are in accordance with other estimates of additive genetic variance derived from animal-model based studies (e.g. 30%: Postma, 2014; 10– 30%: Wood et al., 2016; 22%: Moore et al., 2019). Overall, this result confirms our prediction that the effect of an individual’s own genes on their own phenotype should be larger than the effect of the genotype of a single social partner, even for traits that are often considered highly plastic in response to conspecifics, such as behaviours (Bailey et al., 2018; West-Eberhard, 2003).

### IGEs are largest for behaviours and reproductive traits

IGEs are larger for behaviours and reproductive traits (*social h* ^2^ = 0.06 and 0.05, respectively), compared to other traits categories such as morphology, development, survival, metabolism and physiology (*social h* ^2^ = 0 to 0.02). This higher susceptibility may occur as behavioural and reproductive traits likely involve feedback dynamics between interacting partners, such as mutual escalation of aggression in contests (McGlothlin et al., 2010; Trubenová et al., 2015). Similarly, reproductive traits such as lay date or offspring production should heavily depend on the influence of the mating partner as the reproductive outcome is in part a joint phenotype, contributed to by both parents. Further, behavioural traits often have relatively low estimates of *h*^2^ (Dochtermann et al., 2015; Moore et al., 2019; van Oers & Sinn, 2014) and so our analyses shows that traits with higher *social h*^2^ are those that often have low direct *h*^2^. Therefore, while behaviours may show limited response to direct selection, due to the prominence of IGEs they may respond more to social, group-level, and other types of selection. Altogether, these findings suggest that behaviours and reproductive traits might follow a substantially different evolutionary pathway (including indirect effects), and that evolution for these traits is expected to diverge most from that predicted by DGEs alone.

### Other sources of heterogeneity in IGEs

The additional moderators that we tested to try to explain the high heterogeneity observed showed varying levels of importance. Although life stage explained a non-negligible amount of heteregenity across effect sizes, the magnitude of IGEs did not clearly differ between early and late life stages, which could be because our meta-analysis did not include maternal genetic effects, a type of IGEs expected to be strong for juvenile traits (Moore et al., 2019). The sex of the studied individuals explained a high amount of heterogeneity across effect sizes, with IGEs being most important in female only estimates rather than in those including both sexes. However, we note that we did not have enough male-only estimates for a meaningful comparison (7 effect sizes from only 2 studies) and that 30 out of the 40 female-only estimates corresponded to captive chickens, hence more diversity in study systems is needed to directly compare sexes and confirm this trend. Indeed, considering sex differences in the magnitude of IGEs in future studies will be useful as the sexes often differ in their patterns of social interaction and competition, which is associated with sexually dimorphic traits (Fairbairn et al., 2007; Shine, 1989).

The scarcity of wild studies hampered our assessment of whether IGEs estimates differ compared to studies on captive animals. Fortunately, logistical challenges such as capturing individuals and monitoring their social interactions, along with large sample sizes required by quantitative genetics models (Charmantier et al., 2014), are gradually being met in recent years. The development of large collaborative datasets (e.g. Culina et al. 2021), refined tools for spatial and social network analyses (e.g. Fisher et al. 2019; Radersma 2021), tracking technology (e.g. Nathan et al. 2022), as well as the ability to estimate genetic variances from genomic rather than pedigree data (e.g. Gienapp et al. 2017, Johnston et al. 2022), are expected to lead to more studies estimating IGEs in the wild. These findings mirror results of a meta-analysis estimating DGEs, where there was no support for heritabilities differing between laboratory and field conditions, nor among wild, domestic, or semi-domestic species (Dochtermann et al., 2019), although previous studies show that behaviours measured in the field tended to exhibit higher heritabilities (Mousseau & Roff, 1987; van Oers & Sinn, 2014; Stirling et al., 2002).

### Conclusions

Our meta-analysis across 21 animal species highlights the small but statistically significant relative magnitude of IGEs, yet also highlights their potentially large contribution to adaptive potential. IGEs were most prominent for behavioural and reproduction traits, possibly suggesting a different evolutionary route for these traits, despite the considerable variation across studies. Not accounting for IGEs underestimates the total amount of genetic variance available to selection in most contexts. IGEs can instead enhance the evolutionary potential of traits and speed up evolutionary rates by facilitating a response to selection, possibly promoting the adaptation of populations to rapidly changing environmental conditions. The increase in adaptive potential was mostly due to the relationship between the contribution of IGEs to τ^2^ and group size, but also due to the correlation between IGEs and DGEs tending to be, on average, positive across studies. However, it may take the action of multilevel selection to realise a population’s full potential to respond to selection in the presence of IGEs, and therefore understanding the interplay between IGEs, multilevel selection, and relatedness will be key for accurately predicting evolutionary change. Additionally, future studies should expand our knowledge to underrepresented taxonomic groups, particularly focusing on wild populations, which still remain severely understudied.

## Author contributions

FS: Conceptualization; Methodology; Investigation; Data Curation; Writing – original draft preparation; Writing – review and editing; Project administration

MM: Conceptualization; Methodology; Validation; Formal analysis; Investigation; Writing – original draft preparation; Writing – review and editing; Visualization

AST: Conceptualization; Methodology; Formal analysis; Investigation; Data Curation; Writing – original draft preparation; Writing – review and editing; Visualization

DF: Conceptualization; Methodology; Validation; Formal analysis; Investigation; Writing – original draft preparation; Writing – review and editing; Visualization

## Acknowledgements

We would like to thank Denis Reale, Clint Kelly, and Pierre-Olivier Montiglio for their feedback in the earlier stages of the project, and Daniel W.A. Noble for statistical advice. We thank the participants of the 2023 WAMBAM conference and Alastair Wilson for useful discussion. We also thank Allen Moore, Joel Pick, and one anonymous reviewer for their constructive feedback, which improved the manuscript. We are grateful to the following authors who provided additional data for the meta-analysis when we contacted them: Alastair Wilson, Anasuya Chakrabarty, Cristina Sartori, Esther Ellen, Chang Han, Jane Reid, João Costa e Silva, Jennifer Morinay, Mark Adams, Miriam Piles, Moha Ragab, Ingrid David, Paolo Carnier, Ryan Germain, Setegn Alemu, Simon Evans, and Céline Teplitsky.

## Data and code availability statement

All data and code are available at the following GitHub repository (https://github.com/ASanchez-Tojar/meta-analysis_IGEs). Upon acceptance, all data and code will be provided with a doi via Zenodo.

