## Supplementary Material for "Indirect genetic effects increase the heritable variation available to selection and are largest for behaviours: a meta-analysis"

**SUPPLEMENTARY TEXT**

**Publication bias**

To test for evidence of publication bias, we ran three uni-moderator phylogenetic multilevel meta-regressions with social *h*^2^ (k = 146 effect sizes, N = 40 studies) as the effect size of interest: (1) one including the effect size standard error (SE) to test for evidence of small-study effects (Nakagawa et al. 2022), (2) one including the effect size sampling variance instead of SE, which provides a less downwardly biased estimate for the adjusted meta-analytic mean (Nakagawa et al. 2022), and (3) another including year of publication (mean-centred; Sánchez-Tójar et al. 2018; Nakagawa et al. 2022) to test for evidence of decline effects (also known as time-lag bias). The random effect structure of these three meta-regressions was the same as reported for all models (see main text).

Overall, we found no evidence of publication bias in the primary literature. Regarding small-study effects, social *h*^2^ estimates did not clearly tend to become smaller as precision increased (slope_SE_ = -0.068, 95% CI [-1.077, 0.940]) and SE as a moderator only explained 0.1% of the heterogeneity (*R^2^_marginal_* = 0.001). Since the intercept of that meta-regression was statistically significantly different from zero (intercept = 0.033, 95% CI [0.003, 0.063]), we reran this model but using sampling variance as the moderator instead of SE to obtain a less potentially downwardly biased intercept (i.e. adjusted meta-analytic mean; Nakagawa et al. 2022). This model led to almost an identical estimate for the intercept (intercept = 0.032, 95% CI [0.010, 0.055]), confirming little evidence and impact of small-study effects on our overall estimate of social *h*^2^. In addition, our results also showed little evidence of decline effects (slope_Year of publication (mean-centred)_ = -0.001, 95% CI [-0.005, 0.002]; Figure S6) and the moderator year of publication (mean-centred) only explained 1.8% of the heterogeneity (*R^2^_marginal_* = 0.018). In sum, we found negligible evidence of publication bias in our meta-analytic dataset.

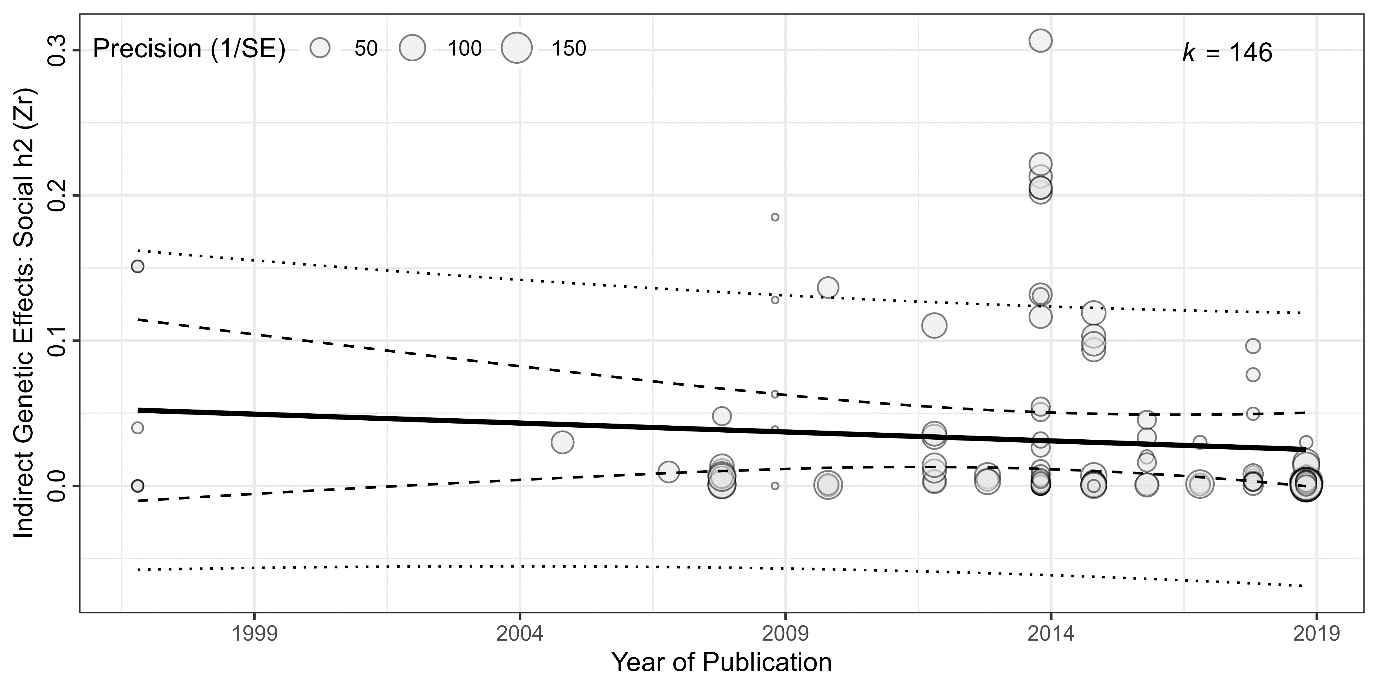

**Figure S6.** Bubble plot showing the relationship between social heritability (*social h*^2^) and year of publication from a phylogenetic multilevel uni-moderator meta-regression with the 95% confidence intervals (dashed lines), 95% prediction intervals (dotted lines) and individual effect sizes (*Zr*) scaled by their precision (semitransparent circles). *k* corresponds to the number of effect sizes.

**Social *h*^2^ (sensitivity analysis)**

Our sensitivity analysis excluding estimates of social *h*^2^ calculated by ourselves and only using estimates reported in the primary studies and supplied by authors showed a very similar social *h*^2^ estimate (*Zr* = 0.042, 95% CI [0.009, 0.076], 95% PI {-0.044, 0.128}, p = 0.015; k = 40 effect sizes, N = 14 studies, 8 species). That is, despite the much smaller dataset, the results are robust both in terms of the meta-analytic mean and its uncertainty.

**Additional analyses**

**Aim 2e: Fixed effect of partner**

Social *h*^2^ was larger when the fixed effect of the partner was not fitted in the animal model (*Zr* = 0.033, 95% CI [0.010, 0.057]; k = 97 effect sizes, N = 32 studies) than when it was (*Zr* = 0.027, 95% CI [-0.009, 0.063]; k = 49 effect sizes, N = 8 studies), but that difference was not statistically significant (*p* = 0.761; Figure S4) and the moderator only explained 0.5% of the heterogeneity (*R^2^_marginal_* = 0.005).

**Aim 2f: Livestock (post-hoc)**

We ran an exploratory meta-regression to test for the difference in social *h^2^* between data obtained from livestock (i.e. *Neovison vison* [k = 6 effect sizes, n = 2 studies], *Sus scrofa* [k = 35 effect sizes, n = 12 studies], *Bos taurus* [k = 1 effect size, n = 1 study], and *Gallus gallus* [k = 28 effect sizes, n = 7 studies]) compared to nonlivestock species. We found that there were no statistically significant differences (p = 0.386) between effect sizes based on livestock species (*Zr* = 0.024 [−0.001, 0.050]; k = 70 effect sizes, N = 20 studies) and those based on non-livestock species (*Zr* = 0.043 [0.010, 0.075]; k = 76 effect sizes, N = 18 studies; *R^2^_marginal_* = 0.037), with the former estimate not differing statistically significantly from zero. Since this meta-regression was run after having seen the results (i.e., post hoc analysis), this finding should be interpreted as explorative (Figure S5).

**Increased sampling variance (sensitivity analysis)**

As in previous meta-analyses of variance-standardised estimates (e.g., Dochtermann et al. 2019, Holtmann et al. 2017), we treated those ICC-type (intra-class correlation) coefficients as Pearson’s correlation coefficients (*r*) throughout our analyses. Although such an analytical approach might be suboptimal, as far as we are aware, there is no better approach to deal with the meta-analysis of such variance-standardised estimates. Treating them as *r* allows us to use the commonly applied Fisher’s Z transformation to calculate *Zr* and its associated variance (*VZr*), which faciliates its meta-analysis. However, such assumption may lead to an underestimation of sampling variance due to, among others, the different levels of replication analyzed in the mixed-effects models from which these estimates are obtained (e.g., due to the structure of the A matrix). To test the robustness of our results to an increase in sampling variance, we decided to rerun our key analyses after having artificially increased sampling variance by a factor of 10 (i.e., *VZr*_new_ = *VZr*_original_ * 10). Such tenfold increase in sampling variance led to virtually the same conclusions throughout, and mostly resulted in slightly broader 95% confidence intervals but slightly narrower 95% prediction intervals, the latter associated with considerable reduction in total heterogeneity (Tenfold model: *𝜎^2^_total_* = 0.0021; *I^2^_total_* = 43.6%; *CV_total_* = 1.11; *M_total_* = 0.63; *VS.* Original model: *𝜎^2^_total_* = 0.0012; *I^2^_total_* = 93.2%; *CV_total_* = 1.46; *M_total_* = 0.70). In addition, the amount of heterogeneity explained by the moderators tended to be larger in the tenfold models, likely also as the consquence of the reduce heterogeneity to be explained. We present all results as figures (S7-11), for more details, see our provided code at: <https://github.com/ASanchez-Tojar/meta-analysis_IGEs>

**Email template sent to authors of papers for data request**

Dear [Corresponding author name],

I hope this email finds you well. I am emailing as I am part of a team collecting data for a meta-analysis to quantify the existence and magnitude of indirect genetic effects, in an attempt to understand their wider importance in evolution. This project is being conducted by myself [Author name and affiliation] along with [Name and affiliation of all other coauthors] and collaborators.

We are aware of the following article for which you are the corresponding author and which contains data we are interested in: [Paper title and citation].

In particular, for the trait [name of traits from the article], we were hoping you could provide us with the following estimates:

•	Trait mean

•	Trait phenotypic variance

•	Unique number of individuals included in the analysis

We are asking for those estimates so that we can calculate parameters such as the ratio of indirect genetic variance to the mean of the trait, and account for sample size in our meta-analysis. In addition to citing your original publication, we would like to add your name in the Acknowledgements section of the eventual publication.

Thanking you very much in advance for your help and looking forward to hearing from you soon.

With best wishes,

[Author name], on behalf of all authors

**SUPPLEMENTARY TABLES**

**Supplementary Table S1.** Meta-analytic means and 95% CI (on the diagonal) for the multiple moderator levels of Aim 2 and the meta-regression (post-hoc) p-values for each pair-wise comparison (off-diagonal).

| **Aim 2a: Trait category** | | | |  |  |  |  |  |
| --- | --- | --- | --- | --- | --- | --- | --- | --- |
|  | Survival | Reproduction | Morphology | | Metabolism & physiology | Development | | Behaviour |
| Survival | 0.025 | 0.354 | 0.384 | | 0.451 | 0.432 | | 0.155 |
|  | [-0.021, 0.071] |  |  |  |  |  |  |  |
| Reproduction |  | 0.053 | 0.033 | | 0.060 | 0.043 | | 0.750 |
|  |  | [0.014,  0.093] |  |  |  |  |  |  |
| Morphology |  |  | 0.001 | | 0.913 | 0.811 | | 0.004 |
|  |  |  | [-0.027, 0.029] | |  |  |  |  |
| Metabolism & physiology |  |  |  | | 0.003 | 0.942 | | 0.014 |
|  |  |  |  | | [-0.033, 0.039] |  |  |  |
| Development |  |  |  | |  | 0.004  [-0.023, 0.031] | | 0.005 |
| Behaviour |  |  |  | |  |  | | 0.061 |
|  |  |  |  | |  |  | | [0.032, 0.091] |

| **Aim 2b: Age** | | | |
| --- | --- | --- | --- |
|  | Juvenile | Both | Adults |
| Juvenile | 0.013 | 0.026 | 0.403 |
|  | [-0.029, 0.055] |  |  |
| Both |  | 0.091 | 0.049 |
|  |  | [0.023, 0.160] |  |
| Adults |  |  | 0.028 |
|  |  |  | [-0.005, 0.062] |

| **Aim 2c: Sex** | | | |
| --- | --- | --- | --- |
|  | Both | Female | Male |
| Both | 0.006 | < 0.001 | 0.412 |
|  | [-0.011, 0.023] |  |  |
| Female |  | 0.063 | 0.524 |
|  |  | [0.041, 0.084] |  |
| Males |  |  | 0.038 |
|  |  |  | [-0.036, 0.112] |

| **Aim 2d: Population type** | | | |
| --- | --- | --- | --- |
|  | Captive | Semi-captive | Wild |
| Captive | 0.030 | 0.750 | 0.495 |
|  | [0.006, 0.053] |  |  |
| Semi-captive |  | 0.020 | 0.448 |
|  |  | [−0.037, 0.076] |  |
| Wild |  |  | 0.048 |
|  |  |  | [−0.000, 0.097] |

| **Aim 2e: Fixed effect of partner (additional analysis)** | | |
| --- | --- | --- |
|  | Partner fixed effect of the partner was not fitted | Partner fixed effect of the partner was fitted |
| Partner fixed effect of the partner was not fitted | 0.033 | 0.761 |
|  | [0.010, 0.057] |  |
| Partner fixed effect of the partner was fitted |  | 0.027 |
|  |  | [-0.009, 0.063] |

**SUPPLEMENTARY FIGURES**

**Figure S1.** Decision tree used to peform the title-and-abstract screening of a systematic literature review on indirect genetic effects.

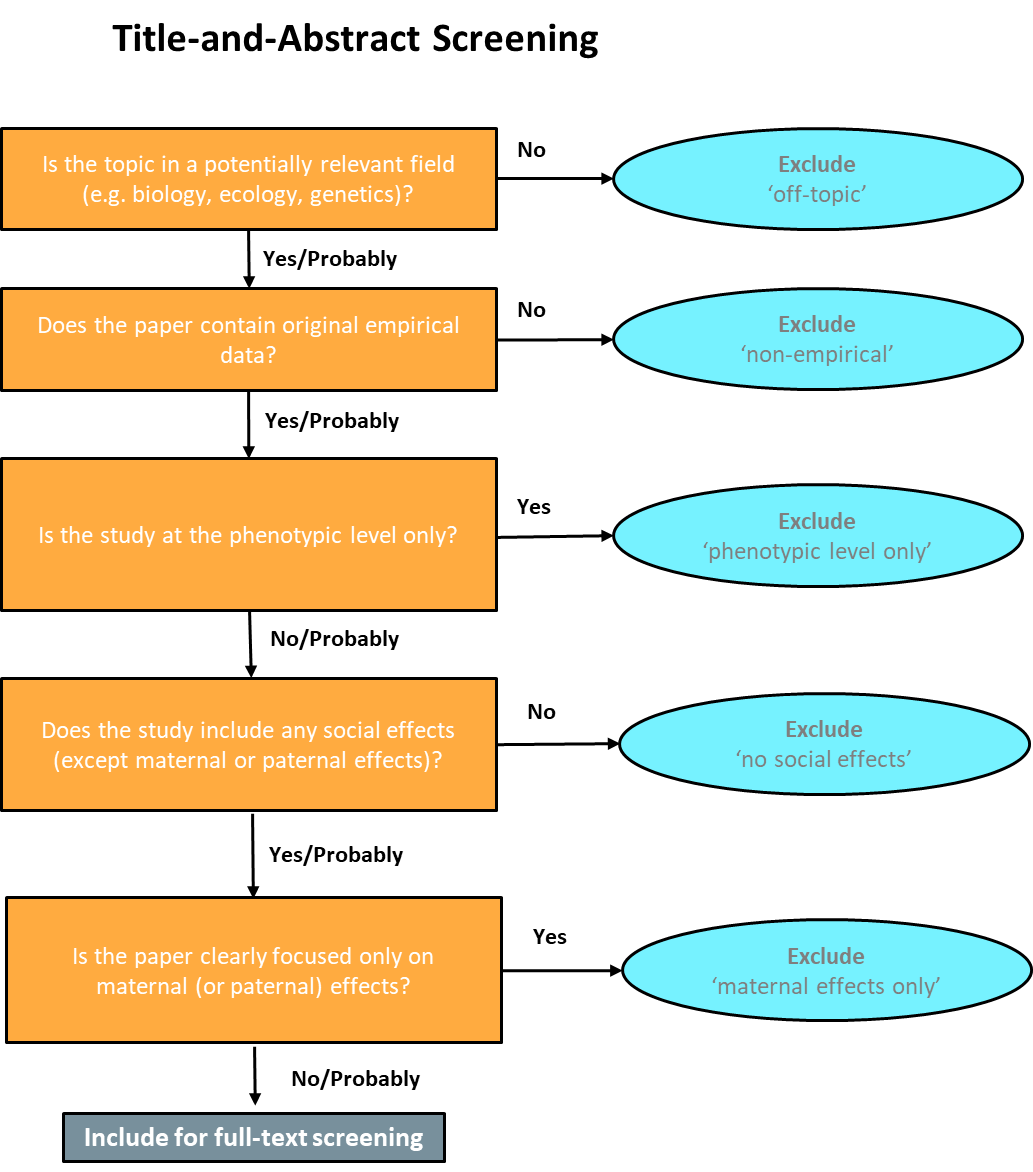

**Figure S2.** Decision tree used to performed the full-text screening of a systematic literature review on indirect genetic effects.

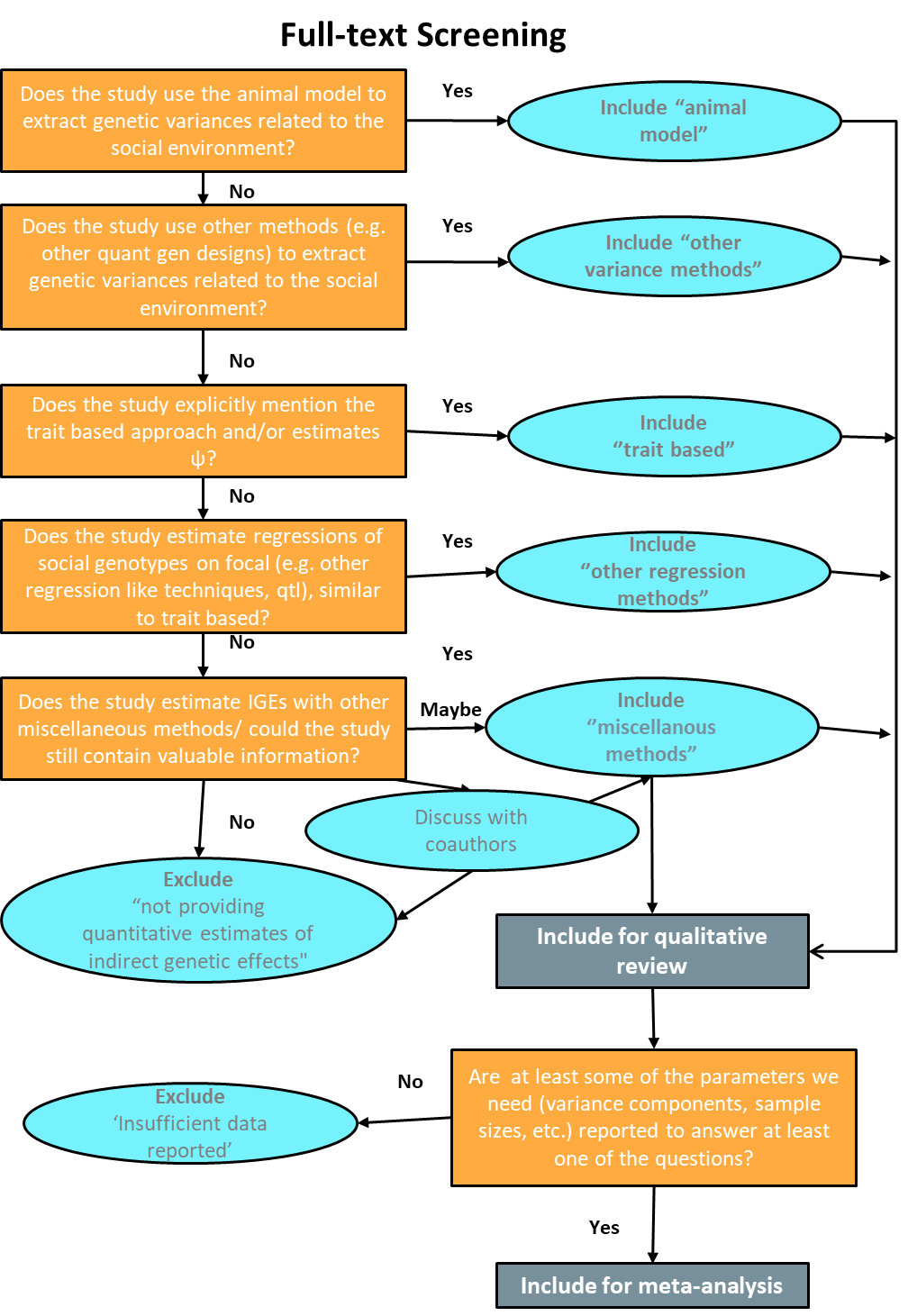

**Figure S3.** PRISMA diagram detailing the process of a systematic literature review on indirect genetic effects.

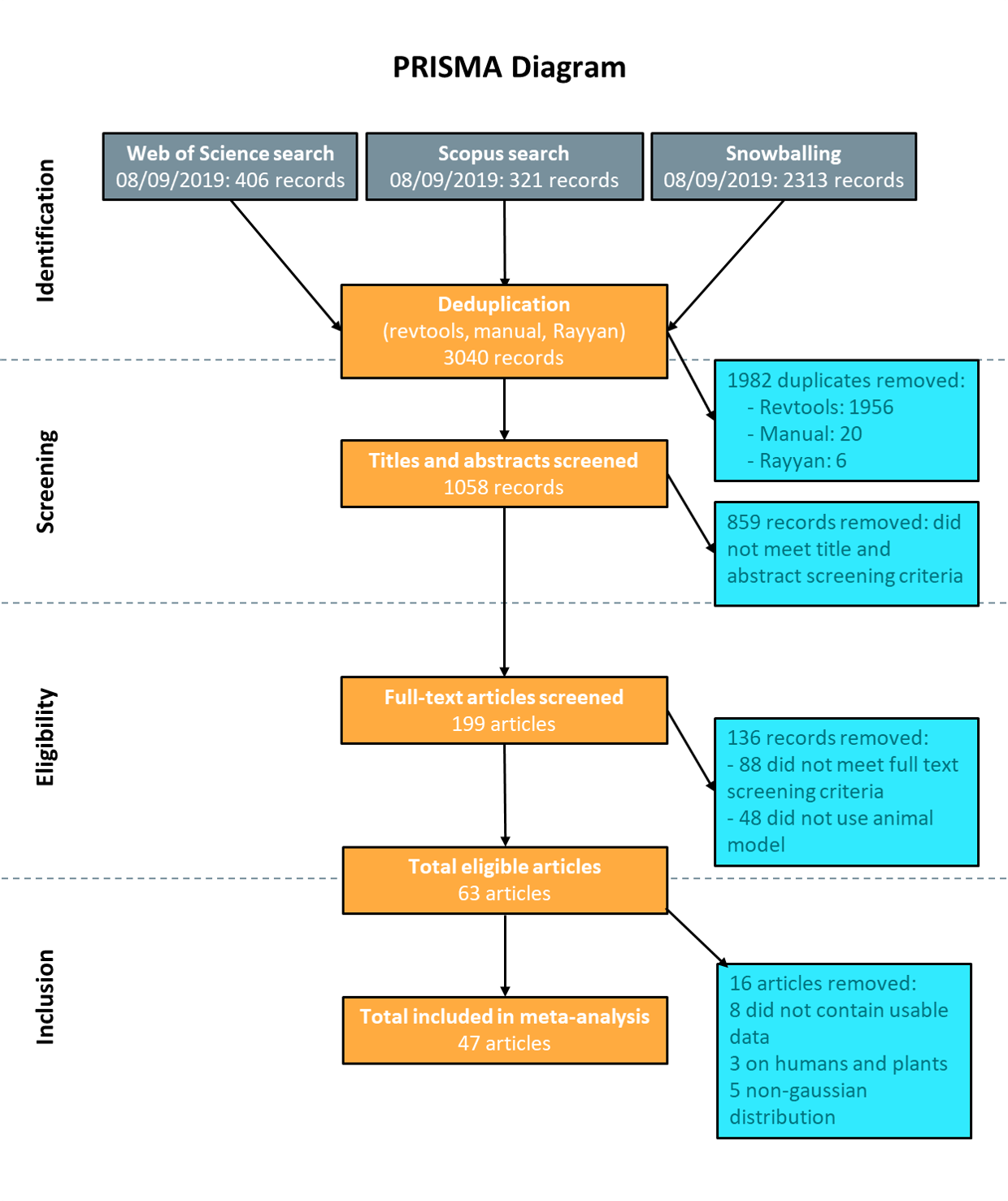

**Figure S4.** Social heritability (*social h*^2^) meta-analytic estimates from phylogenetic multilevel uni-moderator meta-regressions testing the effect of including fixed effects pertaining to social partners in the original animal models (aim 2e, additional analysis). Orchard plot showing the back-transformed meta-analytic mean as *r*, 95% confidence intervals (thick whisker), 95% prediction intervals (thin whisker) and individual effect sizes (*r*) scaled by their precision (semitransparent circles). *k* corresponds to the number of effect sizes, the number of studies is shown in brackets. Note that uncertainty measures (i.e., 95% CI and 95% PI) are generated assuming a normal distribution around the meta-analytic mean, explaining why some intervals may overlap zero substantially despite the data analyzed being bounded between 0 and 1.

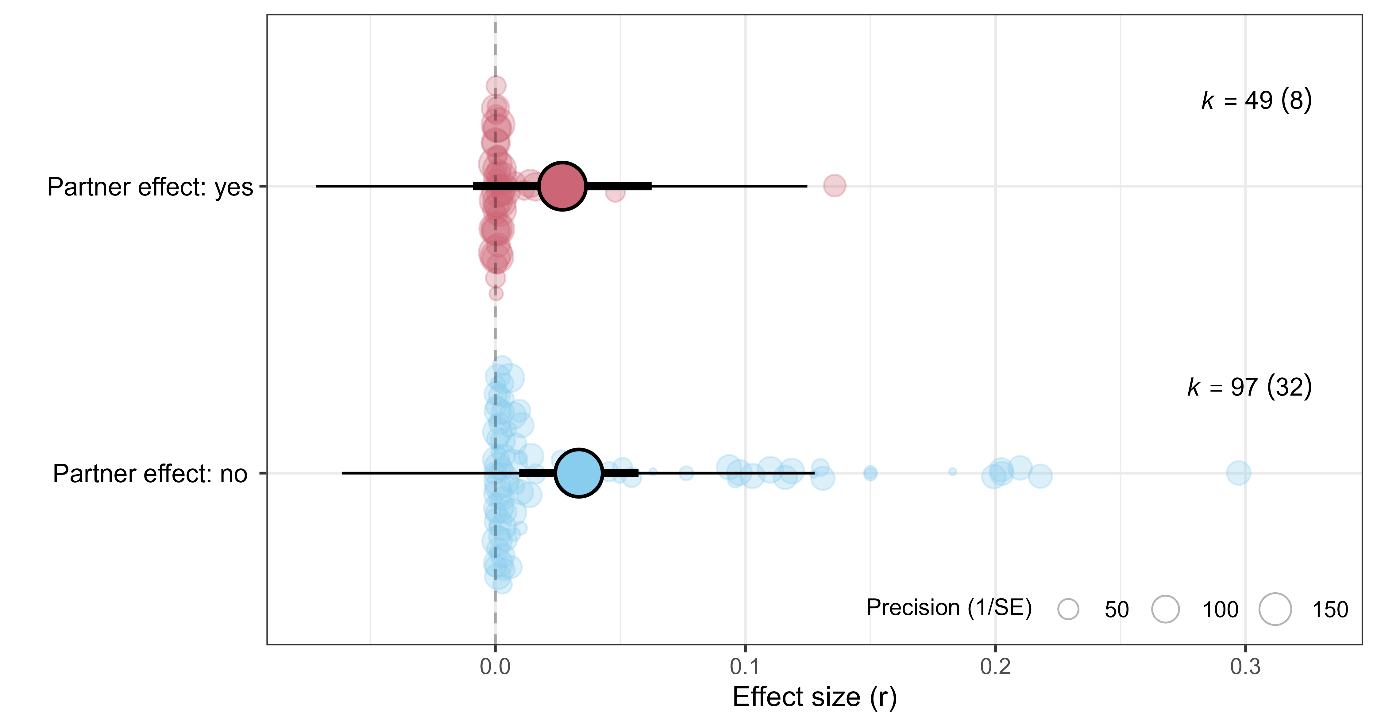

**Figure S5.** Social heritability (*social h*^2^) meta-analytic estimates from phylogenetic multilevel uni-moderator meta-regressions testing the effect of the moderator livestock (aim 2f, post-hoc additional analysis). Orchard plot showing the back-transformed meta-analytic mean as *r*, 95% confidence intervals (thick whisker), 95% prediction intervals (thin whisker) and individual effect sizes (*r*) scaled by their precision (semitransparent circles). *k* corresponds to the number of effect sizes, the number of studies is shown in brackets. Note that uncertainty measures (i.e., 95% CI and 95% PI) are generated assuming a normal distribution around the meta-analytic mean, explaining why some intervals may overlap zero substantially despite the data analyzed being bounded between 0 and 1.

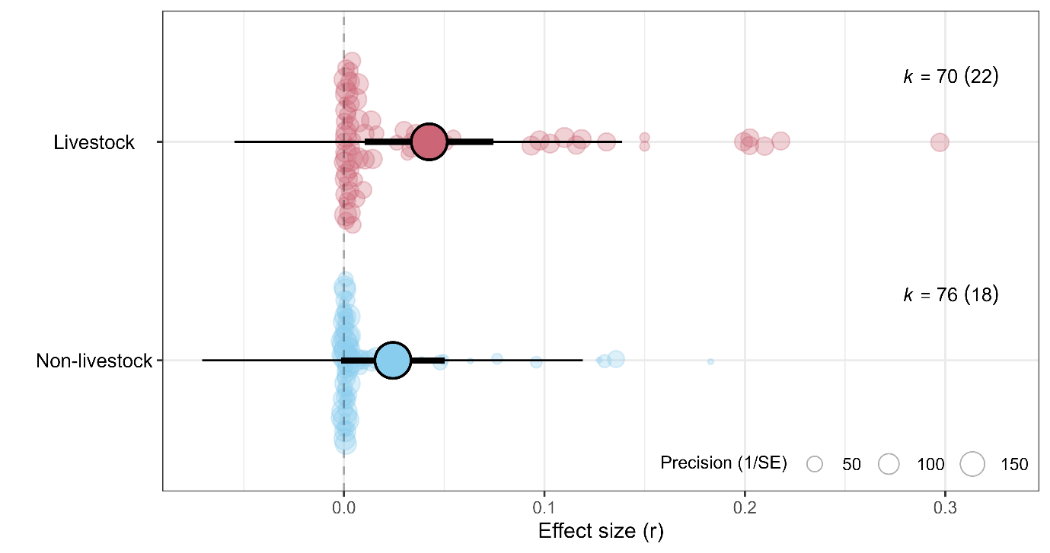

**Figure S7.** Social heritability (*social* *h^2^*) from a phylogenetic multilevel meta-analysis with 40 studies and 146 effect sizes (Aim 1), with a tenfold artificial increase in sampling variance (*VZr*). Orchard plot showing the back-transformed meta-analytic mean as r, 95% confidence intervals (thick whisker), 95% prediction intervals (thin whisker) and individual effect sizes (*r*) scaled by their precision (semitransparent circles). *k* corresponds to the number of effect sizes, the number of studies is shown in brackets. Note that uncertainty measures (i.e., 95% CI and 95% PI) are generated assuming a normal distribution around the meta-analytic mean, explaining why some intervals may overlap zero substantially despite the data analyzed being bounded between 0 and 1. See Figure 2 in the main manuscript to explore the tenfold artificial increase in sampling variance (*VZr*).

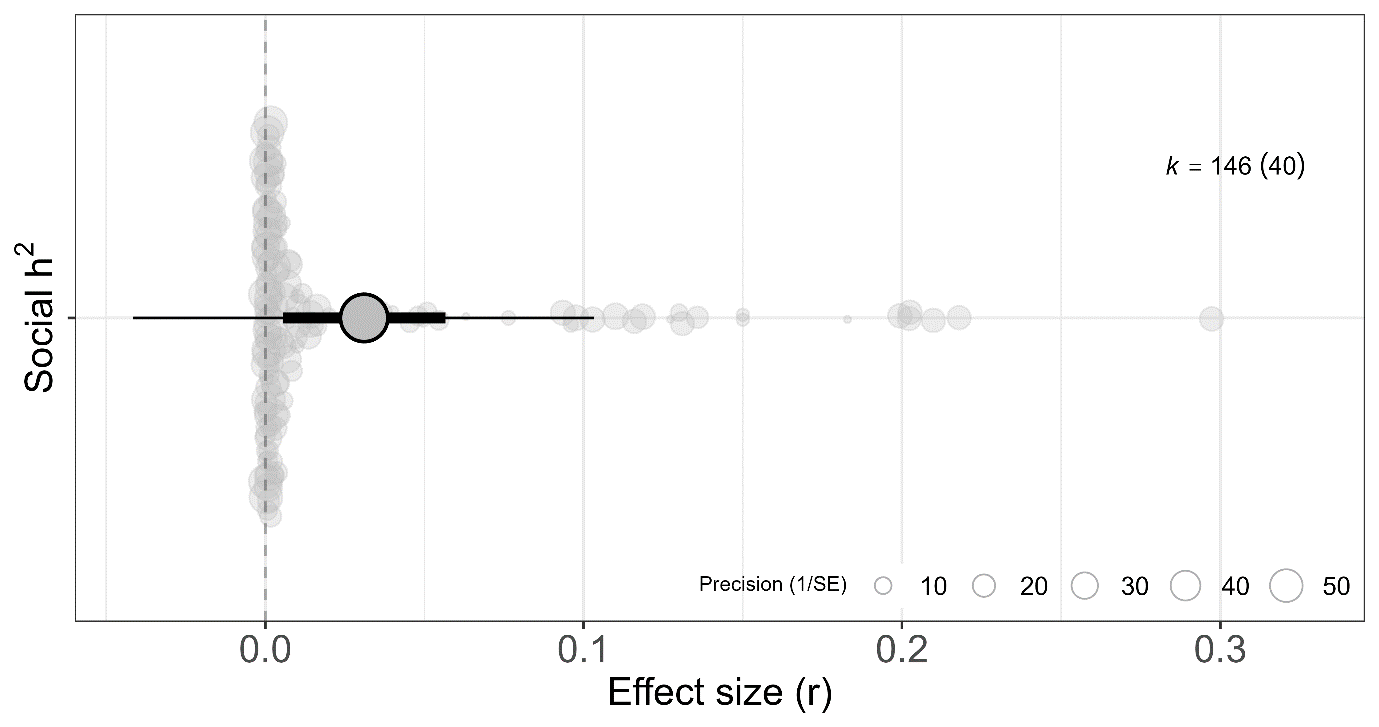

**Figure S8.** Social heritability (*social h*^2^) meta-analytic estimates from a phylogenetic multilevel uni-moderator meta-regression testing the moderator ‘Trait category’ (Aim 2; *R^2^_marginal_* = 43.3%), with a tenfold artificial increase in sampling variance (*VZr*). Orchard plot showing the back-transformed meta-analytic mean as *r*, 95% confidence intervals (thick whisker), 95% prediction intervals (thin whisker) and individual effect sizes (*r*) scaled by their precision (semitransparent circles). *k* corresponds to the number of effect sizes, the number of studies is shown in brackets. See Figure 3a in the main manuscript to explore the tenfold artificial increase in sampling variance (*VZr*).

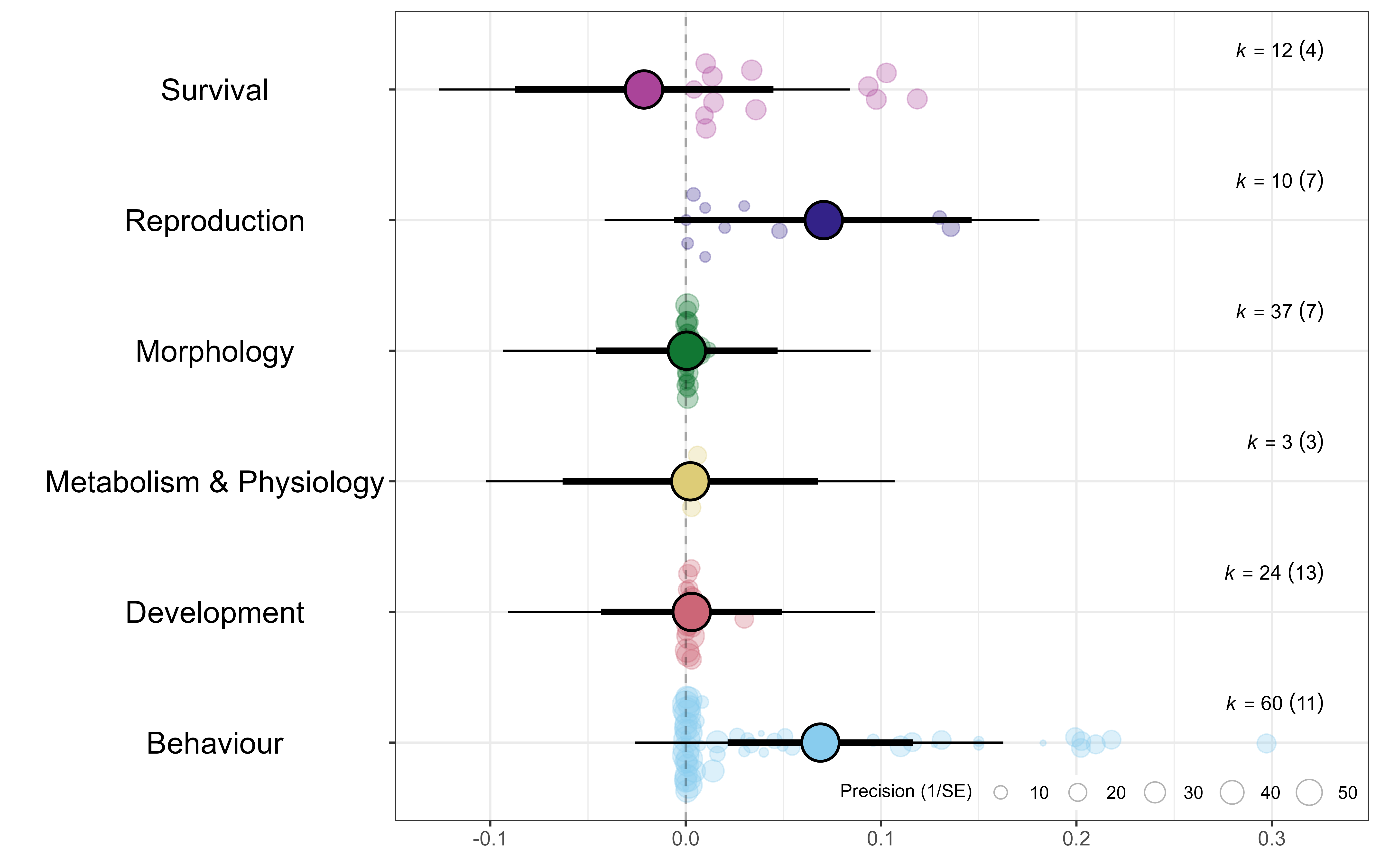

**Figure S9.** Social heritability (*social h*^2^) meta-analytic estimates from a phylogenetic multilevel uni-moderator meta-regression testing the moderator ‘Age’ (Aim 2; *R^2^_marginal_* = 13.8%), with a tenfold artificial increase in sampling variance (*VZr*). Orchard plot showing the back-transformed meta-analytic mean as *r*, 95% confidence intervals (thick whisker), 95% prediction intervals (thin whisker) and individual effect sizes (*r*) scaled by their precision (semitransparent circles). *k* corresponds to the number of effect sizes, the number of studies is shown in brackets. See Figure 3b in the main manuscript to explore the tenfold artificial increase in sampling variance (*VZr*).

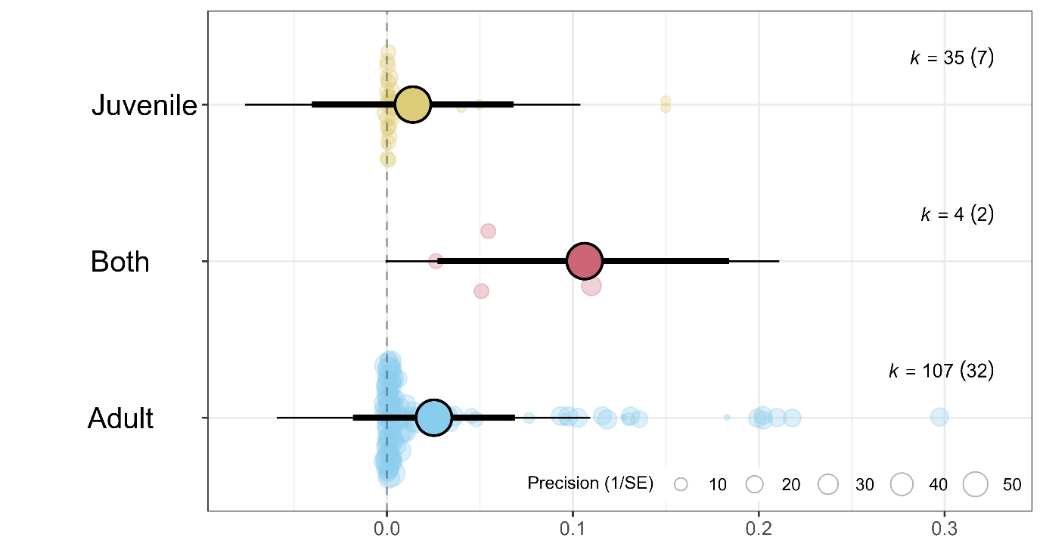

**Figure S10.** IGEs and DGEs meta-analytic estimates from a phylogenetic multilevel uni-moderator meta-regression (Aim 3) comparing narrow-sense *h*^2^ vs. *social h*^2^ (*R^2^_marginal_* = 29.9%), with a tenfold artificial increase in sampling variance (*VZr*). Orchard plot showing the back-transformed meta-analytic mean as *r*, 95% confidence intervals (thick whisker), 95% prediction intervals (thin whisker) and individual effect sizes (*r*) scaled by their precision (semitransparent circles). *k* corresponds to the number of effect sizes, the number of studies is shown in brackets. See Figure 4b in the main manuscript to explore the tenfold artificial increase in sampling variance (*VZr*).

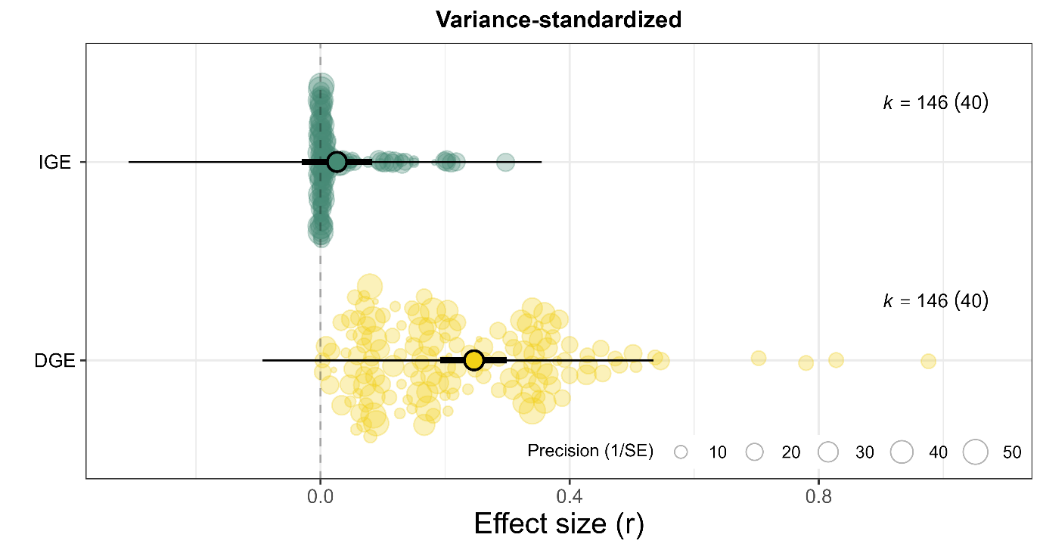

**Figure S11.** Meta-analytic estimates for Aim 4, with a tenfold artificial increase in sampling variance (*VZr*): (a) phylogenetic multilevel meta-regression comparing narrow-sense heritability (*h^2^*) to total heritable variance (τ^2^; *R^2^_marginal_* = 7.0%); (b) phylogenetic multilevel meta-analysis of *r_DGE-IGE_*. Orchard plot showing the original (a) and back-transformed meta-analytic mean as *r*, 95% confidence intervals (thick whisker), 95% prediction intervals (thin whisker) and individual effect sizes (*r*) scaled by their precision (semitransparent circles). *k* corresponds to the number of effect sizes, the number of studies is shown in brackets. See Figure 5 in the main manuscript to explore the tenfold artificial increase in sampling variance (*VZr*).

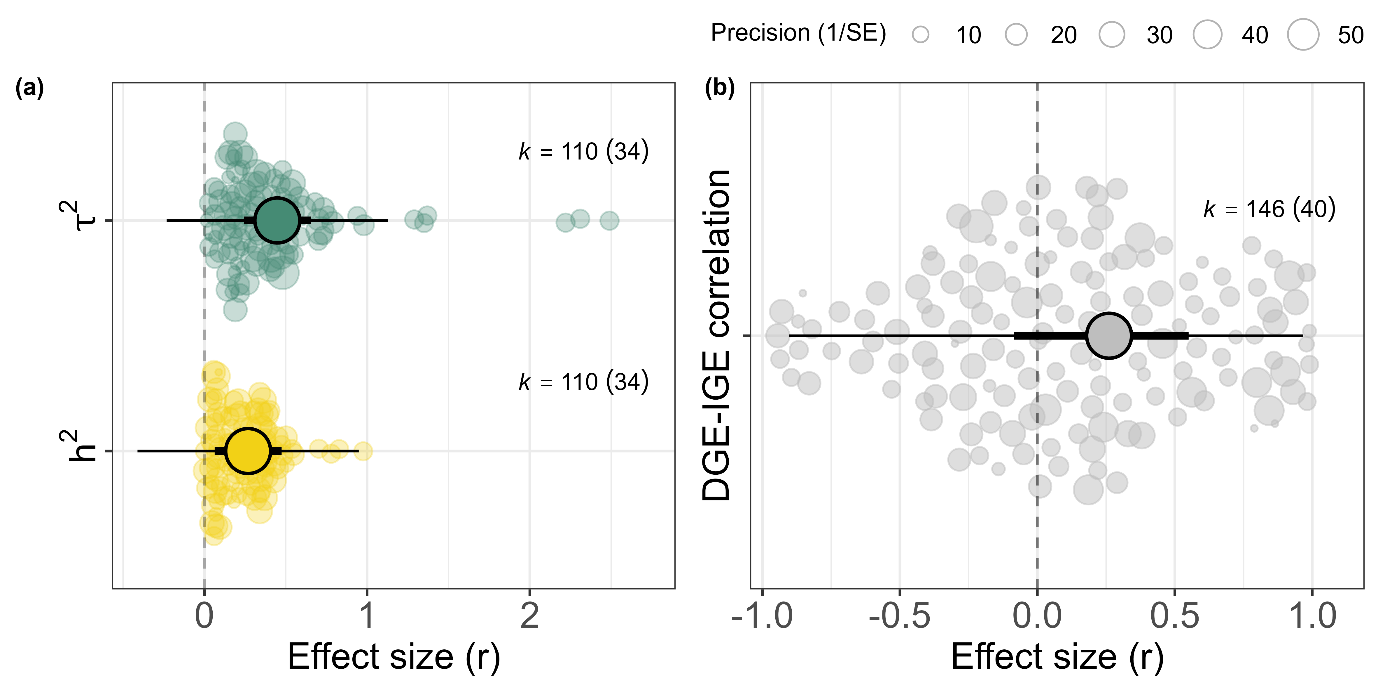
